# Eight thousand years of natural selection in Europe

**DOI:** 10.1101/016477

**Authors:** Iain Mathieson, Iosif Lazaridis, Nadin Rohland, Swapan Mallick, Nick Patterson, Songül Alpaslan Roodenberg, Eadaoin Harney, Kristin Stewardson, Daniel Fernandes, Mario Novak, Kendra Sirak, Cristina Gamba, Eppie R. Jones, Bastien Llamas, Stanislav Dryomov, Joseph Pickrell, Juan Luís Arsuaga, José María Bermúdez de Castro, Eudald Carbonell, Fokke Gerritsen, Aleksandr Khokhlov, Pavel Kuznetsov, Marina Lozano, Harald Meller, Oleg Mochalov, Vayacheslav Moiseyev, Manuel A. Rojo Guerra, Jacob Roodenberg, Josep Maria Vergès, Johannes Krause, Alan Cooper, Kurt W. Alt, Dorcas Brown, David Anthony, Carles Lalueza-Fox, Wolfgang Haak, Ron Pinhasi, David Reich

**Affiliations:** Department of Genetics, Harvard Medical School, Boston, Massachusetts 02115, USA; Broad Institute of MIT and Harvard, Cambridge Massachusetts 02142, USA; Howard Hughes Medical Institute, Harvard Medical School, Boston, Massachusetts 02115, USA; Independent researcher, Santpoort-Noord, The Netherlands; School of Archaeology and Earth Institute, Belfield, University College Dublin, Dublin 4, Ireland; Institute for Anthropological Research, Zagreb 10000, Croatia; Department of Anthropology, Emory University, Atlanta, Georgia, USA; Current address: Centre for GeoGenetics, Natural History Museum of Denmark, University of Copenhagen, Øster Voldgade 5-7, 1350 Copenhagen, Denmark; Smurfit Institute of Genetics, Trinity College Dublin, Dublin 2, Ireland; Australian Centre for Ancient DNA, School of Earth and Environmental Sciences & Environment Institute, University of Adelaide, Adelaide, South Australia 5005, Australia; Laboratory of Human Molecular Genetics, Institute of Molecular and Cellular Biology, Siberian Branch of the Russian Academy of Sciences, Novosibirsk 630090, Russia; Department of Paleolithic Archaeology, Institute of Archaeology and Ethnography, Siberian Branch of the Russian Academy of Sciences, Novosibirsk 630090, Russia; Current Address: New York Genome Center, New York NY, USA; Centro Mixto UCM-ISCIII de Evolución y Comportamiento Humanos, Madrid, Spain.; Departamento de Paleontología, Facultad Ciencias Geológicas, Universidad Complutense de Madrid, Spain.; Centro Nacional de Investigacíon sobre Evolución Humana (CENIEH), 09002 Burgos, Spain; IPHES. Institut Catalá de Paleoecologia Humana i Evolució Social, Campus Sescelades-URV, 43007. Tarragona, Spain.; Area de Prehistoria, Universitat Rovira i Virgili (URV), 43002 Tarragona, Spain.; Netherlands Institute in Turkey, Istiklal Caddesi, Nur-i Ziya Sokak 5, Beyoğlu, Istanbul, Turkey; Volga State Academy of Social Sciences and Humanities, Samara 443099, Russia; State Office for Heritage Management and Archaeology Saxony-Anhalt and State Museum of Prehistory, D-06114 Halle, Germany; Peter the Great Museum of Anthropology and Ethnography (Kunstkamera) RAS, St Petersburg, 199034, Russia; Department of Prehistory and Archaeology, University of Valladolid, Spain; The Netherlands Institute for the Near East, Leiden, RA-2300, The Netherlands; Max Planck Institute for the Science of Human History, D-07745 Jena, Germany; Institute for Archaeological Sciences, University of Tübingen, D-72070 Tübingen, Germany; Danube Private University, A-3500 Krems, Austria; Institute for Prehistory and Archaeological Science, University of Basel, CH-4003 Basel, Switzerland; Institute of Anthropology, Johannes Gutenberg University of Mainz, D-55128 Mainz, Germany; Anthropology Department, Hartwick College, Oneonta, New York 13820, USA; Institute of Evolutionary Biology (CSIC-Universitat Pompeu Fabra), Barcelona, Spain

## Abstract

The arrival of farming in Europe around 8,500 years ago necessitated adaptation to new environments, pathogens, diets, and social organizations. While indirect evidence of adaptation can be detected in patterns of genetic variation in present-day people, ancient DNA makes it possible to witness selection directly by analyzing samples from populations before, during and after adaptation events. Here we report the first genome-wide scan for selection using ancient DNA, capitalizing on the largest genome-wide dataset yet assembled: 230 West Eurasians dating to between 6500 and 1000 BCE, including 163 with newly reported data. The new samples include the first genome-wide data from the Anatolian Neolithic culture, who we show were members of the population that was the source of Europe’s first farmers, and whose genetic material we extracted by focusing on the DNA-rich petrous bone. We identify genome-wide significant signatures of selection at loci associated with diet, pigmentation and immunity, and two independent episodes of selection on height.

---

Natural selection has left its mark on patterns of variation in our genomes^1^, but these patterns are echoes of past events, which are difficult to date and interpret, and are often confounded by neutral processes. Ancient DNA provides a more direct view, and should be a transformative technology for studies of selection just as it has transformed studies of history. Until now, however, the large sample sizes required to detect selection have meant that ancient DNA studies have concentrated on characterizing effects at parts of the genome already believed to have been affected by selection^2-5^.

We assembled genome-wide data from 230 ancient individuals who lived in West Eurasia from 6500 to 1000 BCE (Table 1, Fig. 1a, Supplementary Data Table 1, Supplementary Information section 1). To obtain this dataset, we combined published data from 67 samples from relevant periods and cultures^4-6^, with 163 samples for which we report new data, of which 83 have never previously been analyzed (the remaining 80 samples include 67 whose targeted single nucleotide polymorphism (SNP) coverage we triple from 390k to 1240k^7^; and 13 with shotgun data whose data quality we increase using our enrichment strategy^3,8^). The 163 samples for which we report new data are drawn from 270 distinct individuals who we screened for evidence of authentic DNA^7^. We used in-solution hybridization with synthesized oligonucleotide probes to enrich promising libraries for more than 1.2 million SNPs (“1240k capture”, Methods). The targeted sites include nearly all SNPs on the Affymetrix Human Origins and Illumina 610-Quad arrays, 49,711 SNPs on chromosome X and 32,681 on chromosome Y, and 47,384 SNPs with evidence of functional importance. We merged libraries from the same individual and filtered out samples with low coverage or evidence of contamination to obtain the final set of individuals. The advantage of 1240k capture is that it gives access to genome-wide data from ancient samples with small fractions of human DNA and increases efficiency by targeting sites in the human genome that will actually be analyzed. The effectiveness of the approach can be seen by comparing our results to the largest previously published ancient DNA study, which used a shotgun sequencing strategy^5^. Our median coverage on analyzed SNPs is ~4-times higher even while the mean number of reads generated per sample is 36-times lower (Extended Data Fig. 1).

**Figure 1:**
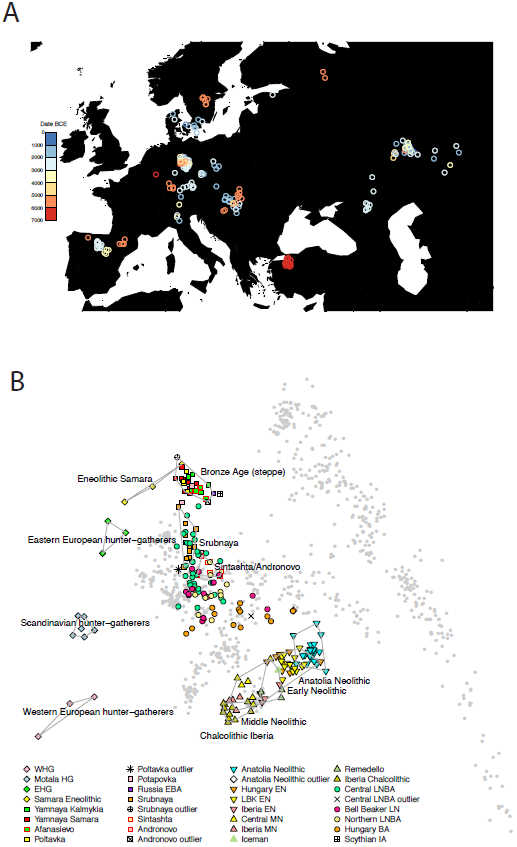
Population relationships of samples. **A:** Locations color-coded by date, with a random jitter added for visibility (8 Afanasievo and Andronovo samples lie further east and are not shown). **B:** Principal component analysis of 777 modern West Eurasian samples (grey), with 221 ancient samples projected onto the first two principal component axes and labeled by culture. **Abbreviations:** [E/M/L]N Early/Middle/Late Neolithic, LBK *Linearbandkeramik,* [E/W]HG Eastern/Western hunter-gatherer, [E]BA [Early] Bronze Age, IA Iron Age.

To learn about the history of archaeological cultures for which genome-wide data is reported for the first time here, we studied either 1,055,209 autosomal SNPs when analyzing 230 ancient individuals alone, or 592,169 SNPs when co-analyzing them with 2,345 present-day individuals genotyped on the Human Origins array^4^. We removed 13 samples either as outliers in ancestry relative to others of the same archaeologically determined culture, or first-degree relatives (Supplementary Data Table 1).

Our sample of 26 Anatolian Neolithic individuals represents the first genome-wide ancient DNA data from the eastern Mediterranean. Our success at analyzing such a large number of samples is likely due to the fact that at the Barcın site–the source of 21 of the working samples–we sampled from the cochlea of the petrous bone^9^, which has been shown to increase the amount of DNA obtained by up to two orders of magnitude relative to teeth (the next-most-promising tissue)^3^. Principal component (PCA) and ADMIXTURE^10^ analysis, shows that the Anatolian Neolithic samples do not resemble any present-day Near Eastern populations but are shifted towards Europe, clustering with Neolithic European farmers (EEF) from Germany, Hungary, and Spain^7^ (Fig. 1b, Extended Data Fig. 2). Further evidence that the Anatolian Neolithic and EEF were related comes from the high frequency (47%; n=15) of Y-chromosome haplogroup G2a typical of ancient EEF samples^7^ (Supplementary Data Table 1), and the low F_ST_ (0.005-0.016) between Neolithic Anatolians and EEF (Supplementary Data Table 2). These results support the hypothesis^7^ of a common ancestral population of EEF prior to their dispersal along distinct inland/central European and coastal/Mediterranean routes. The EEF are slightly more shifted to Europe in the PCA than are the Anatolian Neolithic (Fig. 1b) and have significantly more admixture from Western hunter-gatherers (WHG), shown by *f*4-statistics (|Z|>6 standard errors from 0) and negative *f*3-statistics (|Z|>4)^11^ (Extended Data Table 3). We estimate that the EEF have 7-11% more WHG admixture than their Anatolian relatives (Extended Data Fig. 2, Supplementary Information section 2).

**Figure 2:**
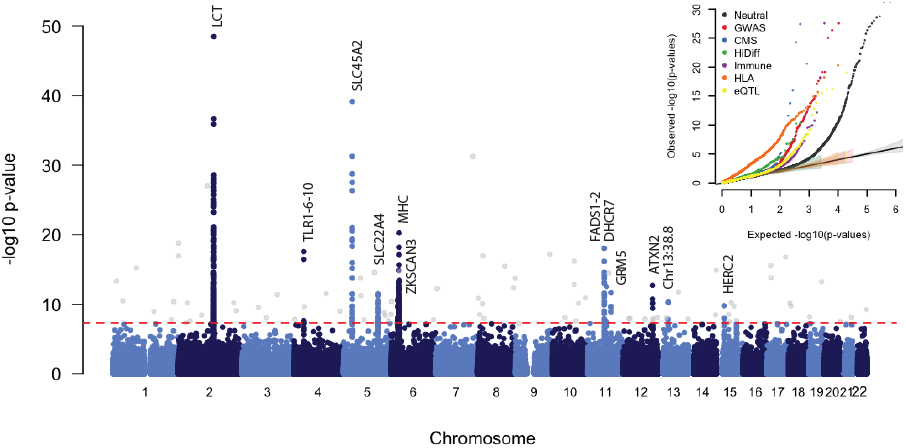
Genome-wide scan for selection. GC-corrected –log_10_ p-value for each marker. The red dashed line represents a genome-wide significance level of 0.5 × 10^−8^. Genome-wide significant points filtered because there were fewer than two other genome-wide significant points within 1Mb are shown in grey. Inset: QQ plots for corrected −log_10_ p-values for different categories of potentially functional SNPs (Methods). Truncated at −log_10_(p-value)=30. All curves are significantly different from neutral expectation.

The Iberian Chalcolithic individuals from El Mirador cave are genetically similar to the Middle Neolithic Iberians who preceded them (Fig. 1b; Extended Data Fig. 2), and have more WHG ancestry than their Early Neolithic predecessors^7^ (|*Z*|>10) (Extended Data Table 3). However, they do not have a significantly different proportion of WHG ancestry (we estimate 23-28%) than the Middle Neolithic Iberians (Extended Data Fig. 2). Chalcolithic Iberians have no evidence of steppe ancestry (Fig. 1b, Extended Data Fig. 2), in contrast to central Europeans of the same period^5,7^. Thus, the “Ancient North Eurasian”-related ancestry that is ubiquitous across present-day Europe^4,7^ arrived in Iberia later than in Central Europe (Supplementary Information section 2).

To understand population transformations in the Eurasian steppe, we analyzed a time transect of 37 samples from the Samara region spanning ~5600-300 BCE and including the Eastern Hunter-gatherer (EHG), Eneolithic, Yamnaya, Poltavka, Potapovka and Srubnaya cultures. Admixture between populations of Near Eastern ancestry and the EHG^7^ began as early as the Eneolithic (5200-4000 BCE), with some individuals resembling EHG and some resembling Yamnaya (Fig. 1b; Extended Data Fig. 2). The Yamnaya from Samara and Kalmykia, the Afanasievo people from the Altai (3300-3000 BCE), and the Poltavka Middle Bronze Age (2900-2200 BCE) population that followed the Yamnaya in Samara, are all genetically homogeneous, forming a tight “Bronze Age steppe” cluster in PCA (Fig. 1b), sharing predominantly R1b Y-chromosomes^5,7^ (Supplementary Data Table 1), and having 48-58% ancestry from an Armenian-like Near Eastern source (Extended Data Table 3) without additional Anatolian Neolithic or Early European Farmer (EEF) ancestry^7^ (Extended Data Fig. 2). After the Poltavka period, population change occurred in Samara: the Late Bronze Age Srubnaya have ~17% Anatolian Neolithic or EEF ancestry (Extended Data Fig. 2). Previous work documented that such ancestry appeared east of the Urals beginning at least by the time of the Sintashta culture, and suggested that it reflected an eastward migration from the Corded Ware peoples of central Europe^5^. However, the fact that the Srubnaya also harbored such ancestry indicates that the Anatolian Neolithic or EEF ancestry could have come into the steppe from a more eastern source. Further evidence that migrations originating as far west as central Europe may not have had an important impact on the Late Bronze Age steppe comes from the fact that the Srubnaya possess exclusively (n=6) R1a Y-chromosomes (Extended Data Table 1), and four of them (and one Poltavka male) belonged to haplogroup R1a-Z93 which is common in central/south Asians^12^, very rare in present-day Europeans, and absent in all ancient central Europeans studied to date.

To study selection, we created a dataset of 1,084,781 autosomal SNPs in 617 samples by merging 213 ancient samples with genome-wide sequencing data from four populations of European ancestry from the 1,000 Genomes Project^13^. Most present-day Europeans can be modeled as a mixture of three ancient populations related to Mesolithic hunter-gatherers (WHG), early farmers (EEF) and steppe pastoralists (Yamnaya)^4,7^, and so to scan for selection, we divided our samples into three groups based on which of these populations they clustered with most closely (Fig. 1b, Extended Data Table 1). We estimated mixture proportions for the present-day European ancestry populations and tested every SNP to evaluate whether its present-day frequencies were consistent with this model. We corrected for test statistic inflation by applying a genomic control correction analogous to that used to correct for population structure in genome-wide association studies. Of ~1 million non-monomorphic autosomal SNPs, the ~50,000 in the set of potentially functional SNPs were significantly more inconsistent with the model than neutral SNPs (Fig. 2), suggesting pervasive selection. Using a conservative significance threshold of p=5.0 × 10^-8^, and a genomic control correction of 1.38, we identified 12 loci that contained at least three SNPs achieving genome-wide significance within 1 Mb of the most associated SNP (Fig. 2, Extended Data Table 2, Extended Data Fig. 3, Supplementary Data Table 3).

**Figure 3:**
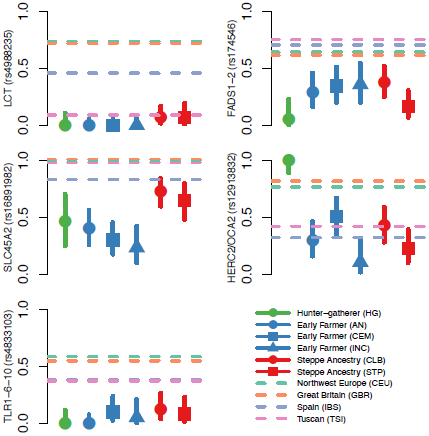
Allele frequencies in different populations. Allele frequencies for five genome-wide significant signals of selection. In each plot, the dots and solid lines show the maximum likelihood frequency estimate and a 1.9-log-likelihood support interval for the derived allele frequency in each ancient population. The four horizontal dashed lines show the allele frequencies in the four modern 1000 Genomes populations. Abbreviations for ancient populations (See Extended Data Table 1); AEN: Anatolian Neolithic; HG: hunter-gatherer; CEM: Central European Early and Middle Neolithic; INC: Iberian Neolithic and Chalcolithic; CLB: Central European Late Neolithic and Bronze Age; STP: Steppe. The Hunter-Gatherer, Early Farmer and Steppe Ancestry classifications correspond approximately to the three populations used in the genome-wide scan with some differences - for example Bell Beakers are included here with CLB but not in the selection scan (See Extended Data Table 1 for details).

The strongest signal of selection is at the SNP (rs4988235) responsible for lactase persistence in Europe^14^. Our data (Fig. 3) strengthens previous reports that an appreciable frequency of lactase persistence in Europe only dates to the last four thousand years^3,5,15^. The allele’s earliest appearance in our data is in a central European Bell Beaker sample (individual I0112) that lived between approximately 2300 and 2200 BCE. Two other independent signals related to diet are located on chromosome 11 near *FADS1* and *DHCR7. FADS1* and *FADS2* are involved in fatty acid metabolism, and variation at this locus is associated with plasma lipid and fatty acid concentration^16^. The selected allele of the most significant SNP (rs174546) is associated with decreased triglyceride levels^16^. Variants at *DHCR7* and *NADSYN1* are associated with circulating vitamin D levels^17^ and our most associated SNP, rs7940244, is highly differentiated across closely related Northern European populations^18^, suggesting selection related to variation in dietary or environmental sources of vitamin D.

Two signals have a potential link to celiac disease. One occurs at the ergothioneine transporter *SLC22A4* that is hypothesized to have experienced a selective sweep to protect against ergothioneine deficiency in agricultural diets^19^. Common variants at this locus are associated with increased risk for ulcerative colitis, celiac disease, and irritable bowel disease and may have hitchhiked to high frequency as a result of this sweep^19-21^. However the specific variant (rs1050152, L503F) that was thought to be the target did not reach high frequency until relatively recently (Extended Data Fig. 4). The signal at *ATXN2/SH2B3*–also associated with celiac disease^20^–shows a similar pattern (Extended Data Fig. 4).

**Figure 4:**
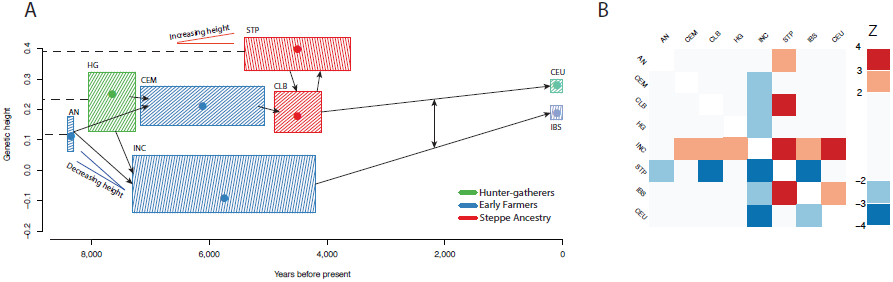
Polygenic selection on height. **A:** Estimated genetic heights for studied populations. Boxes show date ranges and .05 to .95 posterior densities for estimated population mean genetic height (Methods). Dots show the maximum likelihood point estimate of the height. Arrows show major population relationships, with dashed lines representing ancestral populations. Two labeled V’s show our hypothesis for two independent selective events. **B:** Z scores for the polygenic selection scan. The score is positive in each box if the column population is taller than the row population. Abbreviations; AN: Anatolian Neolithic; HG: hunter-gatherer; CEM: Central European Early and Middle Neolithic; INC: Iberian Neolithic and Chalcolithic; CLB: Central European Late Neolithic and Bronze Age; STP: Steppe; CEU: Utah residents with northern and western European ancestry; IBS: Iberian population in Spain.

The second strongest signal in our analysis is at the derived allele of rs16891982 in *SLC45A2,* which contributes to light skin pigmentation and is almost fixed in present-day Europeans but occurred at much lower frequency in ancient populations. In contrast, the derived allele of *SLC24A5* that is the other major determinant of light skin pigmentation in modern Europe appears fixed in the Anatolian Neolithic, suggesting that its rapid increase in frequency to around 0.9 in the Early Neolithic was mostly due to migration (Extended Data Fig. 4). Another pigmentation signal is at *GRM5,* where SNPs are associated with pigmentation possibly through a regulatory effect on nearby TYR^22^. We also find evidence of selection for the derived allele of rs12913832 at *HERC2/OCA2,* which appears to be fixed in Mesolithic hunter-gatherers, and is the primary determinant of blue eye color in present-day Europeans. In contrast to the other loci, the range of frequencies in modern populations is within that of ancient populations (Fig. 3). The frequency increases with higher latitude, suggesting a complex pattern of environmental selection.

The *TLR1-TLR6-TLR10* gene cluster is a known target of selection in Europe^23^, possibly related to resistance to leprosy, tuberculosis or other mycobacteria. There is also a strong signal of selection at the major histocompatibility complex (MHC) on chromosome 6. The strongest signal is at rs2269424 near the genes *PPT2* and *EGFL8* but there are at least six other apparently independent signals in the MHC (Extended Data Fig. 3); and the entire region is significantly more associated than the genome-wide average (residual inflation of 2.07 in the region on chromosome 6 between 29-34 Mb after genome-wide genomic control correction). This could be the result of multiple sweeps, balancing selection, or background selection in this gene-rich region.

We find a surprise in six Scandinavian hunter-gatherers (SHG) from the Motala site in southern Sweden. In three out of six samples, we observe the haplotype carrying the derived allele of rs3827760 in the *EDAR* gene (Extended Data Fig. 5), which affects tooth morphology and hair thickness and has been the subject of a selective sweep in East Asia^24^, and today is at high frequency in East Asians and Native Americans. The *EDAR* derived allele is largely absent in present-day Europe except in Scandinavia, plausibly due to Siberian movements into the region millennia after the date of the Motala samples. The SHG have no evidence of East Asian ancestry^4,7^, suggesting that the *EDAR* derived allele may not have originated not in East Asians as previously suggested^24^. A second surprise is that, unlike closely related western hunter-gatherers, the Motala samples have predominantly derived pigmentation alleles at *SLC45A2* and *SLC24A5.*

We also tested for selection on complex traits. The best-documented example of this process in humans is height, for which the differences between Northern and Southern Europe have driven by selection^25^. To test for this signal in our data, we used a statistic that tests whether trait-affecting alleles are both highly correlated and more differentiated, compared to randomly sampled alleles^26^. We predicted genetic heights for each population and applied the test to all populations together, as well as to pairs of populations (Fig. 4). Using 180 height-associated SNPs^27^ (restricted to 169 where we successfully targeted at least two chromosomes in each population), we detect a significant signal of directional selection on height (p=0.002). Applying this to pairs of populations allows us to detect two independent signals. First, the Iberian Neolithic and Chalcolithic samples show selection for reduced height relative to both the Anatolian Neolithic (p=0.042) and the Central European Early and Middle Neolithic (p=0.003). Second, we detect a signal for increased height in the steppe populations (p=0.030 relative to the Central European Early and Middle Neolithic). These results suggest that the modern South-North gradient in height across Europe is due to both increased steppe ancestry in northern populations, and selection for decreased height in Early Neolithic migrants to southern Europe. We do not observe any other significant signals of polygenetic selection in five other complex traits we tested: body mass index^28^ (p=0.20), waist-to-hip ratio^29^ (p=0.51), type 2 diabetes^30^ (p=0.37), inflammatory bowel disease^21^ (p=0.17) and lipid levels^16^ (p=0.50).

Our results show how ancient DNA can be used to perform a genome-wide scan for selection, and demonstrate selection on loci related to pigmentation, diet and immunity, painting a picture of Neolithic populations adapting to settled agricultural life at high latitudes. For most of the signals we detect, allele frequencies of modern Europeans are outside the range of any ancient populations, indicating that phenotypically, Europeans of four thousand years ago were different in important respects from Europeans today despite having overall similar ancestry. An important direction for future research is to increase the sample size for European selection scans (Extended Data Fig. 6), and to apply this approach to regions beyond Europe and to nonhuman species.

## Acknowledgments

We thank Paul de Bakker, Joachim Burger, Christos Economou, Elin Fornander, Qiaomei Fu, Fredrik Hallgren, Karola Kirsanow, Alissa Mittnik, Iñigo Olalde, Adam Powell, Pontus Skoglund, Shervin Tabrizi, and Arti Tandon for discussions, suggestions about SNPs to include, or contribution to sample preparation or data curation. We thank Svante Pääbo, Matthias Meyer, Qiaomei Fu, and Birgit Nickel for collaboration in developing the 1240k capture reagent. We thank Julio Manuel Vidal Encinas and María Encina Prada for allowing us to resample La Braña 1, and the 1000 Genomes Project for allowing use of the Phase 3 data. I.M. was supported by the Human Frontier Science Program LT001095/2014-L. C.G. was supported by the Irish Research Council for Humanities and Social Sciences (IRCHSS). A.K., P.K. and O.M. were supported by RFBR 𝒩<u>o</u>15-06-01916 and RFHNo15-11-63008 and O.M. by a state grant of the Ministry of education and science of Russia Federation #33.1195.2014/k. J.K. was supported by ERC starting grant APGREID and DFG grant KR 4015/1-1. K.W.A. was supported by DFG grant AL 287 / 14-1. W.H. and B.L. were supported by Australian Research Council DP130102158. R.P. was supported by ERC starting grant ADNABIOARC (263441), and an Irish Research Council ERC support grant. D.R. was supported by U.S. National Science Foundation HOMINID grant BCS-1032255, U.S. National Institutes of Health grant GM100233, and the Howard Hughes Medical Institute.

## Author Information

The aligned sequences are available through the European Nucleotide Archive under accession number [to be made available on publication]. The Human Origins genotype datasets including ancient individuals can be found at (http://genetics.med.harvard.edu/reichlab/Reich_Lab/Datasets.html). Reprints and permissions information is available at www.nature.com/reprints. The authors declare no competing financial interests. Readers are welcome to comment on the online version of the paper. Correspondence and requests for materials should be addressed to I.M.

(iain), W.H., R.P. or D.R.

## Author Contributions

W.H., R.P. and D.R. supervised the study. S.A.R., J.L.A., J.M.B., E.C., F.G., A.K., P.K., M.L., H.M., O.M., V.M., M.A.R., J.R., J.M.V., J.K., A.C., K.W.A., D.B., D.A., C.L., W.H., R.P. and D.R. assembled archaeological material. I.M., I.L., N.R., S.M., N.P., S.D., J.P., W.H. and D.R. analysed genetic data. N.R., E.H., K.S., D.F., M.N., K.S., C.G., E.R.J., B.L., C.L. and W.H. performed wet laboratory ancient DNA work. I.M., I.L. and D.R. wrote the manuscript with help from all co-authors.

## Methods

### Ancient DNA analysis

We screened 433 next generation sequencing libraries from 270 distinct samples for authentic ancient DNA using previously reported protocols^7^. All libraries that we included in nuclear genome analysis were treated with uracil-DNA-glycosylase (UDG) to reduce characteristic errors of ancient DNA^31^.

We performed in-solution enrichment for a targeted set of 1,237,207 SNPs using previously reported protocols^4,7,32^. The targeted SNP set merges 394,577 SNPs first reported in Ref. 7 (390k capture), and 842,630 SNPs first reported in ref.^33^ (840k capture). For 67 samples for which we newly report data in this study, there was pre-existing 390k capture data^7^. For these samples, we only performed 840k capture and merged the resulting sequences with previously generated 390k data. For the remaining samples, we pooled the 390k and 840k reagents together to produce a single enrichment reagent. We attempted to sequence each enriched library up to the point where we estimated that it was economically inefficient to sequence further. Specifically, we iteratively sequenced more and more from each sample and only stopped when we estimated that the expected increase in the number of targeted SNPs hit at least once would be less than one for every 100 new read pairs generated. After sequencing, we filtered out samples with <30,000 targeted SNPs covered at least once, with evidence of contamination based on mitochondrial DNA polymorphism^32^, an appreciable rate of heterozygosity on chromosome X despite being male^34^, or an atypical ratio of X to Y sequences

Of the targeted SNPs, 47,384 are “potentially functional” sites chosen as follows (with some overlap): 1,290 SNPs identified as targets of selection in Europeans by the Composite of Multiple Signals (CMS) test^1^; 21,723 SNPS identified as significant hits by genome-wide association studies, or with known phenotypic effect (GWAS); 1,289 SNPs with extremely differentiated frequencies between HapMap populations^35^ (HiDiff); 9,116 immunochip SNPs chosen for study of immune phenotypes (Immune); 347 SNPs phenotypically relevant to South America (mostly altitude adaptation SNPs in *EGLN1* and *EPAS1),* 5,387 SNPs which tag *HLA* haplotypes (HLA) and 13,672 expression quantitative trait loci^36^ (eQTL).

### Population history analysis

We used two datasets for population history analysis. *“HO”* consists of 592,169 SNPs, taking the intersection of the SNP targets and the Human Origins SNP array^4^; we used this dataset for co-analysis of present-day and ancient samples. *“HOIll”* consists of 1,055,209 SNPs that additionally includes sites from the Illumina genotype array^37^; we used this dataset for analyses only involving the ancient samples.

On the *HO* dataset, we carried out principal components analysis in *smartpca^38^* using a set of 777 West Eurasian individuals^4^, and projected the ancient individuals with the option “lsqproject: YES”. We carried out ADMIXTURE analysis on a set of 2,345 present-day individuals and the ancient samples after pruning for LD in PLINK 1.9 (https://www.cog-genomics.org/plink2)^39^ with parameters “-indep-pairwise 200 25 0.4”. We varied the number of ancestral populations between K=2 and K=20, and used cross-validation (--cv) to identify the value of K=17 to plot in Extended Data Fig. 2f.

We used ADMIXTOOLS^11^ to compute *f*-statistics, determining standard errors with a Block Jackknife and default parameters. We used the option “inbreed: YES” when computing *f*_3_-statistics of the form *f_3_(Ancient;* Ref_1_, Ref_2_) as the *Ancient* samples are represented by randomly sampled alleles rather than by diploid genotypes. For the same reason, we estimated F_ST_ genetic distances between populations on the *HO* dataset with at least two individuals in *smartpca* also using the “inbreed: YES” option.

We estimated ancestral proportions as in Supplementary Information section 9 of Ref. 7, using a method that fits mixture proportions on a *Test* population as a mixture of *N Reference* populations by using *f*_4_-statistics of the form *f*_4_(*Test* or *Ref,* O_1_; O_2_, O_3_) that exploit allele frequency correlations of the *Test* or *Reference* populations with triples of *Outgroup* populations. We used a set of 15 world outgroup populations^4,7^. In Extended Data Fig. 2, we added WHG and EHG as outgroups for those analyses in which they are not used as reference populations.

We determined sex by examining the ratio of aligned reads to the sex chromosomes^40^. We assigned Y-chromosome haplogroups to males using version 9.1.129 of the nomenclature of the International Society of Genetic Genealogy (www.isogg.org), restricting analysis using *samtools*^41^ to sites with map quality and base quality of at least 30, and excluding 2 bases at the ends of each sequenced fragment.

### Genome-wide scan for selection

For most ancient samples, we did not have sufficient coverage to make reliable diploid calls. We therefore used the counts of sequences covering each SNP to compute the likelihood of the allele frequency in each population. Suppose that at a particular site, for each population we have *M* samples with sequence level data, and *N* samples for which we had hard genotype calls (Loschbour, Stuttgart and the 1,000 Genomes samples). For samples *i =* 1... *N,* with genotype data, we observe *X* copies of the reference allele out of 2*N* total chromosomes. For each of samples *i = (N +* 1)… (*N + M),* with sequence level data, we observe *R_i_* sequences with the reference allele out of *T_i_* total sequences. Then, dropping the subscript *i* for brevity, the likelihood of the population reference allele frequency, *p* given data 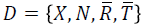 is given by

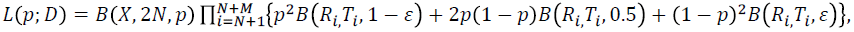
where 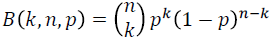 is the binomial probability distribution and *ε* is a small probability of error, which we set to 0.001. We write *ℓ*(*p; D*) for the log-likelihood. To estimate allele frequencies, for example in Fig. 3 or for the polygenic selection test, we maximized this likelihood numerically for each population.

To scan for selection across the genome, we used the following test. Consider a single SNP. Assume that we can model the allele frequencies *p_mod_* in *A* modern populations as a linear combination of allele frequencies in *B* ancient populations *p_anc_*. That is, *p_mod_ = CP_anc_*, where *C* is an *A* by *B* matrix with rows summing to 1. We have data *D_j_* from population *j* which is some combination of sequence counts and genotypes as described above. Then, writing 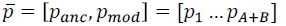 the log-likelihood of the allele frequencies equals the sum of the log-likelihoods for each population.

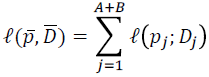

To detect deviations in allele frequency from expectation, we test the null hypothesis *H*_0_*: p_mod_ = C p_anc_* against the alternative *H*_1_:*p_mod_* unconstrained. We numerically maximize this likelihood in both the constrained and unconstrained model and use the fact that twice the difference in log-likelihood is approximately 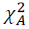 distributed to compute a test statistic and p-value.

We defined the ancient source populations by the “Selection group 1” label in Extended Data Table 1 and Supplementary Table 1 and used the 1000 Genomes CEU, GBR, IBS and TSI as the present-day populations. We removed SNPs that were monomorphic in all four of these modern populations as well as in 1000 Genomes Yoruba (YRI). We do not use FIN as one of the modern populations, because they do not fit this three-population model well. We estimate the proportions of (HG, EF, SA) to be CEU=(0.196, 0.257, 0.547), GBR=(0.362,0.229,0.409), IBS= (0, 0.686, 0.314) and TSI=(0, 0.645, 0.355). In practice we found that there was substantial inflation in the test statistic, most likely due to unmodeled ancestry or additional drift. To address this, we applied a genomic control correction^42^, dividing all the test statistics by a constant, *λ,* chosen so that the median p-value matched the median of the null 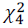 distribution. Excluding sites in the potentially functional set, we estimated *λ =* 1.38 and used this value as a correction throughout. One limitation of this test is that, although it identifies likely signals of selection, it cannot provide much information about the strength or date of selection. If the ancestral populations in the model are, in fact, close to the real ancestral populations, then any selection must have occurred after the first admixture event (in this case, after 6500 BCE), but if the ancestral populations are mis-specified, even this might not be true.

To estimate power, we randomly sampled allele counts from the full dataset, restricting to polymorphic sites with a mean frequency across all populations of <0.1. We then simulated what would happen if the allele had been under selection in all of the modern populations by simulating a Wright-Fisher trajectory with selection for 50, 100 or 200 generations, starting at the observed frequency. We took the final frequency from this simulation, sampled observations to replace the actual observations in that population, and counted the proportion of simulations that gave a genome-wide significant result after GC correction (Extended Data Fig. 6a). We resampled sequence counts for the observed distribution for each population to simulate the effect of increasing sample size, assuming that the coverage and distribution of the sequences remained the same (Extended Data Fig. 6b).

We investigated how the genomic control correction responded when we simulated small amounts of admixture from a highly diverged population (Yoruba; 1000 Genomes YRI) into a randomly chosen modern population. The genomic inflation factor increases from around 1.38 to around 1.51 with 10% admixture, but there is little reduction in power (Extended Fig. 6c). Finally, we investigated how robust the test was to misspecification of the mixture matrix *C*. We reran the power simulations using a matrix *C*′ *= pC +* (1 – *p*)*R* for *p* ∈ [0,1] where *R* was a random matrix chosen so that for each modern population, the mixture proportions of the three ancient populations were jointly normally distributed on [0,1]. Increasing *p* increases the genomic inflation factor and reduces power, demonstrating the advantage of explicitly modeling the ancestries of the modern populations (Extended Fig. 6d).

### Test for polygenic selection

We implemented the test for polygenic selection described by Ref. 26. This evaluates whether trait-associated alleles, weighted by their effect size, are over-dispersed compared to randomly sampled alleles, in the directions associated with the effects measured by genome-wide association studies (GWAS). For each trait, we obtained a list of significant SNP associations and effect estimates from GWAS data, and then applied the test both to all populations combined and to selected pairs of populations. We restricted the list of GWAS associations to 169 SNPs where we observed at least two chromosomes in all tested populations (selection population 2). We estimated frequencies in each population by computing the MLE, using the likelihood described above. For each test, we sampled SNPs frequency matched in 20 bins, computed the test statistic *Q_X_* and for ease of comparison, converted these to Z scores, signed according the direction of the genetic effects. Theoretically *Q_X_* has a *χ*^2^ distribution but in practice, it is over-dispersed. Therefore, we report bootstrap p-values computed by sampling 10,000 sets of frequency matched SNPs.

To estimate population-level genetic height in Fig. 4A, we assumed a uniform prior on [0,1] for the distribution of all height-associated alleles, and then sampled from the posterior joint frequency distribution of the alleles, assuming they were independent, using a Metropolis-Hastings sampler with a N(0,0.001) proposal density. We then multiplied the sampled allele frequencies by the effect sizes to get a distribution of genetic height.

## Extended data

**Extended Data Table 1:** 230 ancient individuals analyzed in this study.

| By population | Population | Date range | N | Out | Rel | Eff N Chr | Selection population 1 | Selection population 2 |
| --- | --- | --- | --- | --- | --- | --- | --- | --- |
|  | WHG | 8.2-8.0 kya | 3 | 0 | 0 | 4.66 | HG | HG |
|  | Motala_HG | 7.9-7.5 kya | 6 | 0 | 0 | 5.19 | HG | HG |
|  | Anatolia_Neolithic | 8.4-8.3 kya | 24 | 1 | 1 | 22.49 | EF | AN |
|  | Hungary_EN | 7.7-7.7 kya | 10 | 0 | 0 | 8.81 | EF | CEM |
|  | LBK_EN | 7.5-7.1 kya | 15 | 0 | 0 | 11.15 | EF | CEM |
|  | Central_MN | 5.9-5.8 kya | 6 | 0 | 0 | 3.66 | EF | CEM |
|  | Iberia_EN | 7.3-7.2 kya | 4 | 0 | 1 | 3.54 | EF | INC |
|  | Iberia_MN | 5.9-5.6 kya | 4 | 0 | 0 | 3.47 | EF | INC |
|  | Iberia_Chalcolithic | 4.8-4.2 kya | 12 | 0 | 2 | 5.93 | EF | INC |
|  | Remedello | 5.5-5.1 kya | 3 | 0 | 0 | 0.93 | EF | - |
|  | Iceman | 5.4-5.1 kya | 1 | 0 | 0 | 1.90 | EF | - |
|  | Central_LNBA | 4.9-4.6 kya | 35 | 1 | 2 | 17.55 | SA | CLB |
|  | Yamnaya_Samara | 5.4-4.9 kya | 9 | 0 | 0 | 6.55 | SA | STP |
|  | Yamnaya_Kalmykia | 5.3-4.7 kya | 6 | 0 | 0 | 3.50 | SA | STP |
|  | Afanasievo | 5.3-5.0 kya | 5 | 0 | 0 | 3.01 | SA | STP |
|  | Poltavka | 4.9-4.7 kya | 4 | 1* | 0 | 4.28 | SA | STP |
|  | Sintashta | 4.3-4.1 kya | 5 | 0 | 0 | 2.35 | SA | STP |
|  | Potapovka | 4.2-4.1 kya | 3 | 0 | 0 | 0.66 | SA | STP |
|  | Srubnaya | 3.9-3.6 kya | 12 | 1* | 1 | 7.68 | SA | STP |
|  | Andronovo | 3.8-3.6 kya | 3 | 1* | 0 | 3.87 | SA | STP |
|  | Russia_EBA | 4.9-4.5 kya | 1 | 0 | 0 | 0.21 | SA | - |
|  | Northern_LNBA | 4.9-4.5 kya | 10 | 0 | 0 | 3.81 | SA | - |
|  | Bell_Beaker_LN | 4.5-4.5 kya | 17 | 0 | 1 | 6.64 | SA^ | CLB |
|  | Hungary_BA | 4.2-4.1 kya | 12 | 0 | 0 | 4.18 | SA^ | CLB |
|  | EHG | 7.7-7.6 kya | 3 | 0 | 0 | 2.15 | - | - |
|  | Samara_Eneolithic | 7.2-6.0 kya | 3 | 0 | 0 | 1.07 | - | - |
|  | Scythian_IA | 2.4-2.2 kya | 1 | 0 | 0 | 1.26 | - | - |

| By selection population | Selection population 1 | Date range | N | Out | Rel | Eff N Chr | Description |
| --- | --- | --- | --- | --- | --- | --- | --- |
|  | EF | 8.4-4.2 kya | 79 | 0 | 0 | 61.88 | Early Farmer |
|  | HG | 8.2-7.5 kya | 9 | 0 | 0 | 9.85 | Hunter-gatherer |
|  | SA | 5.4-3.6 kya | 93 | 3 | 0 | 52.14 | Steppe Ancestry |
| By selection population | Selection population 2 | Date range | N | Out | Rel | Eff N Chr | Description |
|  | AN | 8.4-8.3 kya | 24 | 0 | 0 | 22.49 | Anatolian Neolithic |
|  | CEM | 7.7-5.8 kya | 31 | 0 | 0 | 23.62 | Central European Early and Middle Neolithic |
|  | INC | 7.3-4.2 kya | 20 | 0 | 0 | 12.95 | Iberian Neolithic and Chalcolithic |
|  | HG | 8.2-7.5 kya | 9 | 0 | 0 | 9.85 | Hunter-gatherer |
|  | CLB | 4.9-4.1 kya | 64 | 0 | 0 | 28.38 | Central European Late Neolithic and Bronze Age |
|  | STP | 5.4-3.6 kya | 47 | 3 | 0 | 30.58 | Steppe |

**Population:** samples grouped by a combination of date, source, archaeology and genetics. **Date range:** approximate date range of samples in this group. **N:** Number of individuals sampled. **Out:** Number of PCA outliers (marked with an asterisk if used in selection analysis). **Rel:** Number of related individuals removed. **Eff N Chr:** Average over sites of the effective number of chromosomes when we use genotype likelihoods. Computed as 2 per called site for samples with genotype calls, or $2 - 0.5^{(c-1)}$ for samples with read depth $c$ . **Selection population 1:** Coarse population labels (marked with a caret if not used in genome-wide scan) **Selection population 2:** Fine population labels. **Abbreviations:** [E/M/L]N Early/Middle/Late Neolithic, LBK *Linearbandkeramik*, [E/S/W]HG Eastern/Scandinavian/Western hunter-gatherer, [E]BA [Early] Bronze Age, IA Iron Age.

**Extended Data Table 2:** Twelve genome-wide significant signals of selection.

| Lead SNP | Chromosome | Position (hg19) | P Value | Range (Mb) | Genes | Potential function |
| --- | --- | --- | --- | --- | --- | --- |
| rs4988235 | 2 | 136,608,646 | 3.19E-49 | 135.3-137.3 | MCM6,LCT | Lactase persistence*<sup>14</sup> |
| rs16891982 | 5 | 33,951,693 | 7.05E-40 | 33.8-34.0 | SLC45A2 | Skin pigmentation*<sup>43</sup> |
| rs2269424 | 6 | 32,132,233 | 5.41E-21 | 29.9-33.1 | MHC region | Immunity* |
| rs174546 | 11 | 61,569,830 | 8.18E-19 | 61.5-61.6 | FADS1,FADS2 | Fatty acid metabolism*<sup>16,44,45</sup> |
| rs4833103 | 4 | 38,815,502 | 2.58E-18 | 38.7-38.8 | TLR1,TLR6,TLR10 | Immunity<sup>46,47</sup> |
| rs653178 | 12 | 112,007,756 | 1.96E-13 | 111.9-112.6 | ATXN2,SH2B3 | Unknown |
| rs7944926 | 11 | 71,165,625 | 2.86E-13 | 71.2-71.2 | DHCR7,NADSYN1 | Vitamin D metabolism<sup>17,48</sup> |
| rs7119749 | 11 | 88,515,022 | 2.03E-12 | 88.5-88.9 | GRM5 | Skin pigmentation<sup>22</sup> |
| rs272872 | 5 | 131,675,864 | 2.56E-12 | 131.4-131.8 | SLC22A4 | Ergothioneine transport<sup>49</sup> |
| rs6903823 | 6 | 28,322,296 | 2.96E-11 | 28.3-28.7 | ZKSCAN3,ZSCAN31 | Autophagy<sup>50</sup>, Lung function<sup>51</sup> |
| rs1979866 | 13 | 38,825,900 | 4.60E-11 | 38.1-38.8 | - | Unknown |
| rs12913832 | 11 | 28,365,618 | 1.5E-10 | 29.9-30.1 | HERC2,OCA2 | Eye color*<sup>52,53</sup> |

**Chromosome/Position/Range:** Co-ordinates (hg19) of the SNP with the most significant signal, and the approximate range in which genome-wide significant SNPs are found. **Genes:** Genes in which the top SNP is located, and selected nearby genes. **Potential function:** Function of the gene, or specific trait under selection. Marked with an asterisk if the signal was still genome-wide significant in an analysis that used only the populations that correspond best to the three ancestral populations (WHG, Anatolian Neolithic and Bronze Age steppe), resulting in a less powerful test with the effective number of chromosomes analyzed at the average SNP reduced from 125 to 50, a genomic control correction of 1.32, and five genome-wide significant loci that are a subset of the original twelve.

**Extended Data Table 3:** Key *f*-statistics used to support claims about population history.

| A | B | C | D | $f_4(A, B, C, D)$ | Z | Number of SNPs | Interpretation |
| --- | --- | --- | --- | --- | --- | --- | --- |
| Anatolia_Neolithic | LBK_EN | WHG | Chimp | -0.00114 | -6.8 | 1003751 | Early European Farmers had more WHG ancestry than Anatolian Neolithic |
| Anatolia_Neolithic | Hungary_EN | WHG | Chimp | -0.00212 | -11.9 | 929553 |  |
| Anatolia_Neolithic | Iberia_EN | WHG | Chimp | -0.00244 | -9.6 | 904437 |  |
| Iberia_EN | Iberia_Chalcolithic | WHG | Chimp | -0.00311 | -10.5 | 802471 | Iberian Chalcolithic had more WHG ancestry than Iberian Early Neolithic |
| Iberia_MN | Iberia_Chalcolithic | WHG | Chimp | 0.00010 | 0.3 | 779905 | Iberian Chalcolithic did not have more WHG ancestry than Iberian Middle Neolithic |
| EHG | Samara_Eneolithic | MA1 | Chimp | 0.00140 | 2.3 | 463388 | First dilution of Ancient North Eurasian ancestry (prior to the Bronze Age Yamnaya culture) |
| EHG | Yamnaya_Samara | MA1 | Chimp | 0.00513 | 10.6 | 645211 |  |
| Samara_Eneolithic | Yamnaya_Samara | MA1 | Chimp | 0.00366 | 7.6 | 482492 |  |
| EHG | Yamnaya_Samara | Armenian | Chimp | -0.00191 | -6.1 | 547370 | Contribution of Near Eastern ancestry to the Bronze Age Yamnaya culture |
| EHG | Yamnaya_Kalmykia | Armenian | Chimp | -0.00180 | -5.4 | 536989 |  |
| Samara_Eneolithic | Yamnaya_Samara | Armenian | Chimp | -0.00100 | -3.3 | 405599 |  |
| EHG | Poltavka | Armenian | Chimp | -0.00175 | -4.9 | 541983 |  |
| Yamnaya_Samara | Yamnaya_Kalmykia | MA1 | Chimp | -0.00010 | -0.3 | 675630 | Stability of Ancient North Eurasian ancestry between Early Bronze Age Yamnaya from Kalmykia and Samara, and the Middle Bronze Age Poltavka |
| Yamnaya_Samara | Poltavka | MA1 | Chimp | -0.00014 | -0.4 | 673726 |  |
| Yamnaya_Kalmykia | Poltavka | MA1 | Chimp | 0.00012 | 0.3 | 659346 |  |
| Yamnaya_Samara | Srubnaya | MA1 | Chimp | 0.00151 | 5.1 | 691149 | Second dilution of Ancient North Eurasian ancestry (prior to the Late Bronze Age Srubnaya culture) |
| Yamnaya_Kalmykia | Srubnaya | MA1 | Chimp | 0.00161 | 4.8 | 676735 |  |
| Poltavka | Srubnaya | MA1 | Chimp | 0.00164 | 4.5 | 674756 |  |
| Yamnaya_Samara | Srubnaya | LBK_EN | Chimp | -0.00225 | -11.4 | 974659 | Arrival of Early European Farmer-related ancestry prior to the Late Bronze Age Srubnaya culture. Statistics with Anatolia_Neolithic instead of LBK_EN are similar (Z<-8, not shown). |
| Yamnaya_Kalmykia | Srubnaya | LBK_EN | Chimp | -0.00264 | -11.4 | 951827 |  |
| Poltavka | Srubnaya | LBK_EN | Chimp | -0.00210 | -9.0 | 948968 |  |
| EHG | Yamnaya_Samara | Armenian | LBK_EN | -0.00080 | -5.0 | 559478 | Different source of dilution of Ancient North Eurasian ancestry prior to the Yamnaya (Near Eastern) vs. prior to the Srubnaya (Early European Farmer-related) |
| EHG | Yamnaya_Kalmykia | Armenian | LBK_EN | -0.00086 | -5.2 | 548882 |  |
| EHG | Poltavka | Armenian | LBK_EN | -0.00069 | -4.1 | 553996 |  |
| Yamnaya_Samara | Srubnaya | Armenian | LBK_EN | 0.00138 | 13.1 | 585240 |  |
| Yamnaya_Kalmykia | Srubnaya | Armenian | LBK_EN | 0.00142 | 11.3 | 574333 |  |
| Poltavka | Srubnaya | Armenian | LBK_EN | 0.00134 | 10.7 | 577082 |  |

| Ref<sub>1</sub> | Ref<sub>2</sub> | Test | $f_3(\text{Test; Ref1, Ref2})$ | Z | Number of SNPs | Interpretation |
| --- | --- | --- | --- | --- | --- | --- |
| WHG | Anatolia_Neolithic | Hungary_EN | -0.00412 | -6.7 | 548445 | Early European farmers were formed by admixture between Anatolia Neolithic and WHG (the non-significant signal in the Iberia_EN may be due to genetic drift specific to this population) |
| WHG | Anatolia_Neolithic | LBK_EN | -0.00257 | -4.6 | 654357 |  |
| WHG | Anatolia_Neolithic | Iberia_EN | 0.00179 | 1.4 | 389101 |  |
| EHG | Armenian | Poltavka | -0.00539 | -3.9 | 213055 | Early and Middle Bronze Age steppe pastoralists were formed by admixture between EHG and a population of Near Eastern ancestry |
| EHG | Armenian | Yamnaya_Kalmykia | -0.00537 | -4.2 | 213996 |  |
| EHG | Armenian | Yamnaya_Samara | -0.00586 | -6.2 | 276568 |  |
| LBK_EN | Yamnaya_Samara | Srubnaya | -0.00630 | -11.2 | 584111 | Srubnaya was formed by admixture between populations related to Yamnaya and Early European Farmers |

**Extended Data Figure 1:**
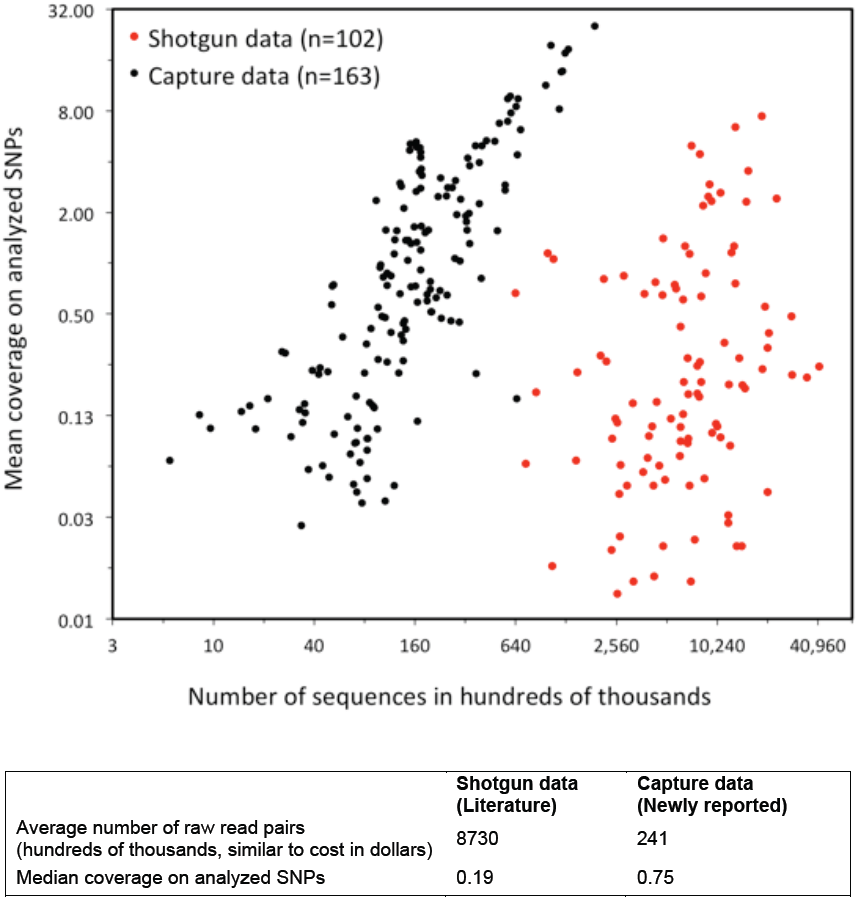
Efficiency and cost-effectiveness of 1240k capture. We plot the number of raw sequences against the mean coverage of analyzed SNPs after removal of duplicates, comparing the 163 samples for which capture data are reported in this study, against the 102 samples analyzed by shotgun sequencing in ref. ^5^ We caution that the true cost is more than that of sequencing alone.

**Extended Data Figure 2:**
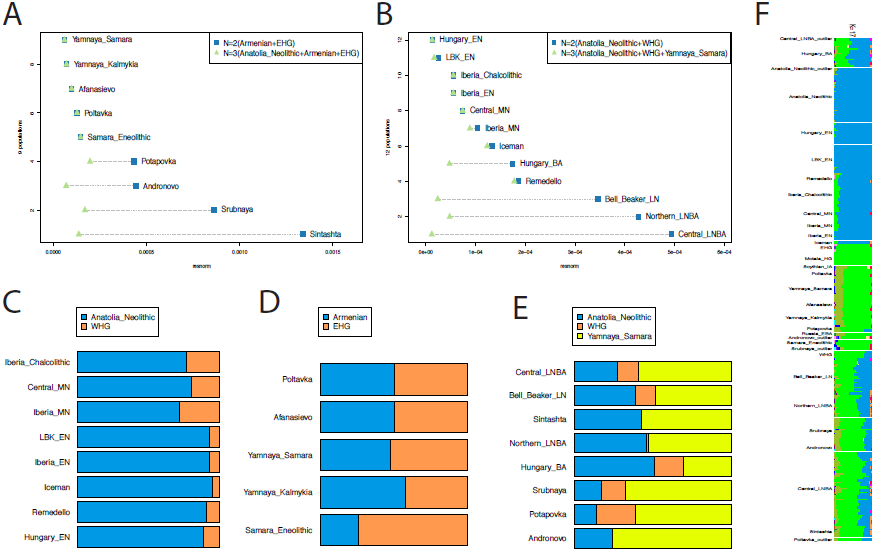
Early isolation and later admixture between farmers and steppe populations. **A:** Mainland European populations later than 3000 BCE are better modeled with steppe ancestry as a 3^rd^ ancestral population. **B:** Later (post-Poltavka) steppe populations are better modeled with Anatolian Neolithic as a 3^rd^ ancestral population. **C:** Estimated mixture proportions of mainland European populations without steppe ancestry. **D:** Estimated mixture proportions of Eurasian steppe populations without Anatolian Neolithic ancestry. **E:** Estimated mixture proportions of later populations with both steppe and Anatolian Neolithic ancestry. **F:** ADMIXTURE plot at k=17 showing population differences over time and space.

**Extended Data Figure 3:**
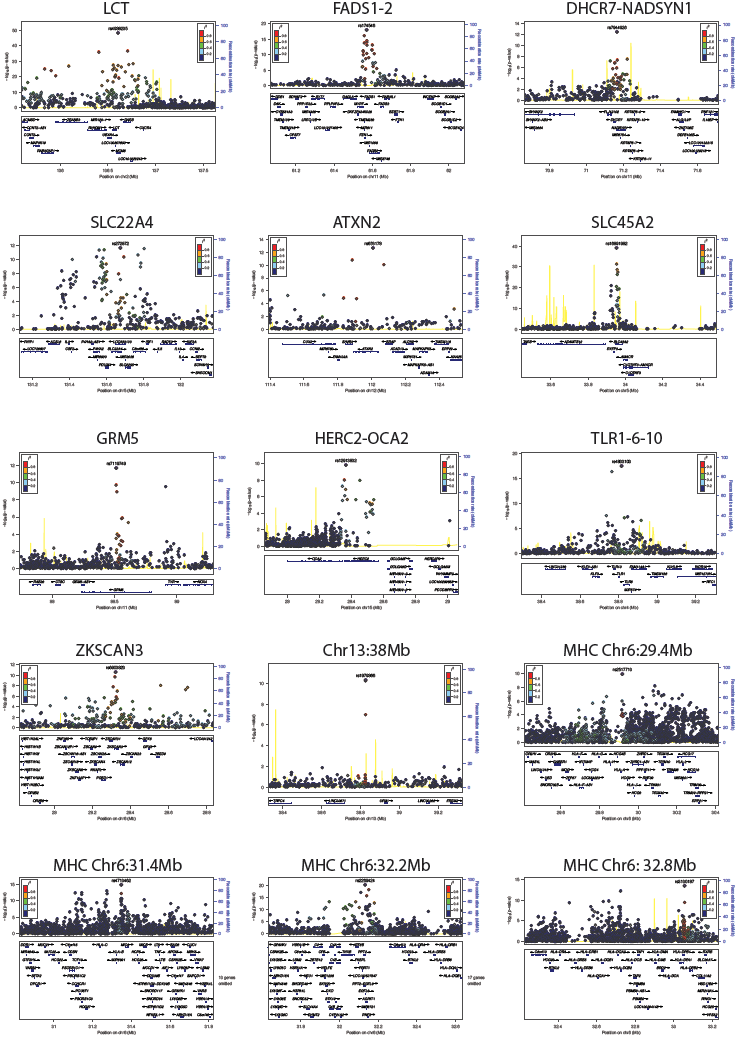
Regional association plots (Locuszoom^54^) for genome-wide significant signals. Points show the –log10 P-value for each SNP, colored according to their LD with the most associated SNP. The blue line shows the recombination rate, with scale on right hand axis. Genes are shown in the lower panel of each subplot.

**Extended Data Figure 4:**
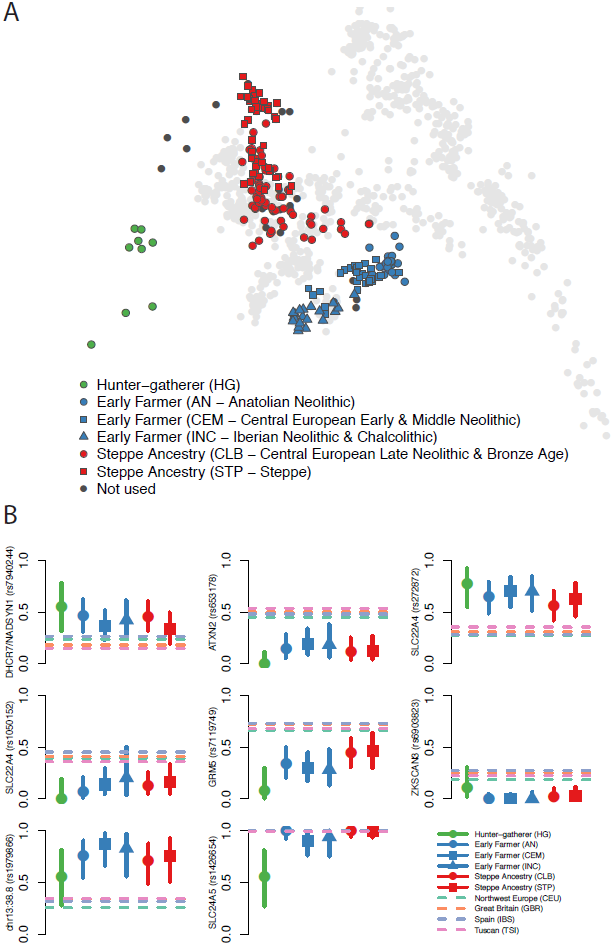
PCA of selection populations and derived allele frequencies for genome-wide significant signals. **A:** Ancient samples projected onto principal components of modern samples, as in Fig. 1, but labeled according to selection populations defined in Extended Table 1. **B:** Allele frequency plots as in Fig. 3. Six signals not included in Fig. 3 – for *SLC22A4* we show both rs272872, which is our strongest signal, and rs1050152, which was previously hypothesized to be under selection – and we also show SLC24A5, which is not genome-wide significant but is discussed in the main text.

**Extended Data Figure 5:**
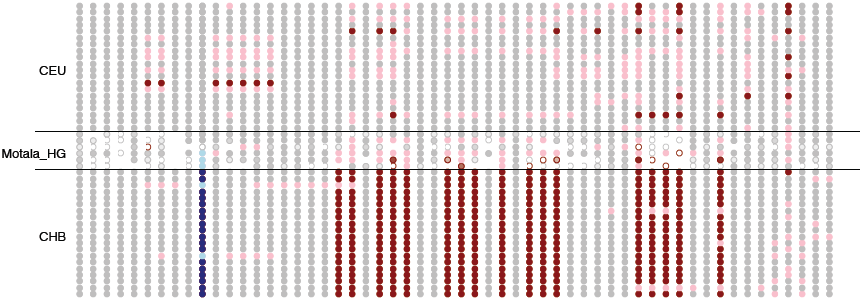
Motala haplotypes carrying the derived, selected EDAR allele. This figure compares the genotypes at all sites within 150kb of rs3827760 (in blue) for the 6 Motala samples and 20 randomly chosen CHB (Chinese from Beijing) and CEU (Utah residents with northern and western European ancestry) samples. Each row is a sample and each column is a SNP. Grey means homozygous for the major (in CEU) allele. Pink denotes heterozygous and red homozygous for the other allele. For the Motala samples, an open circle means that there is only a single sequence otherwise the circle is colored according to the number of sequences observed. Three of the Motala samples are heterozygous for rs3827760 and the derived allele lies on the same haplotype background as in present-day East Asians. The only other ancient samples with evidence of the derived EDAR allele in this dataset are two Afanasievo samples dating to 3300-3000 BCE, and one Scythian dating to 400-200 BCE (not shown).

**Extended Data Figure 6:**
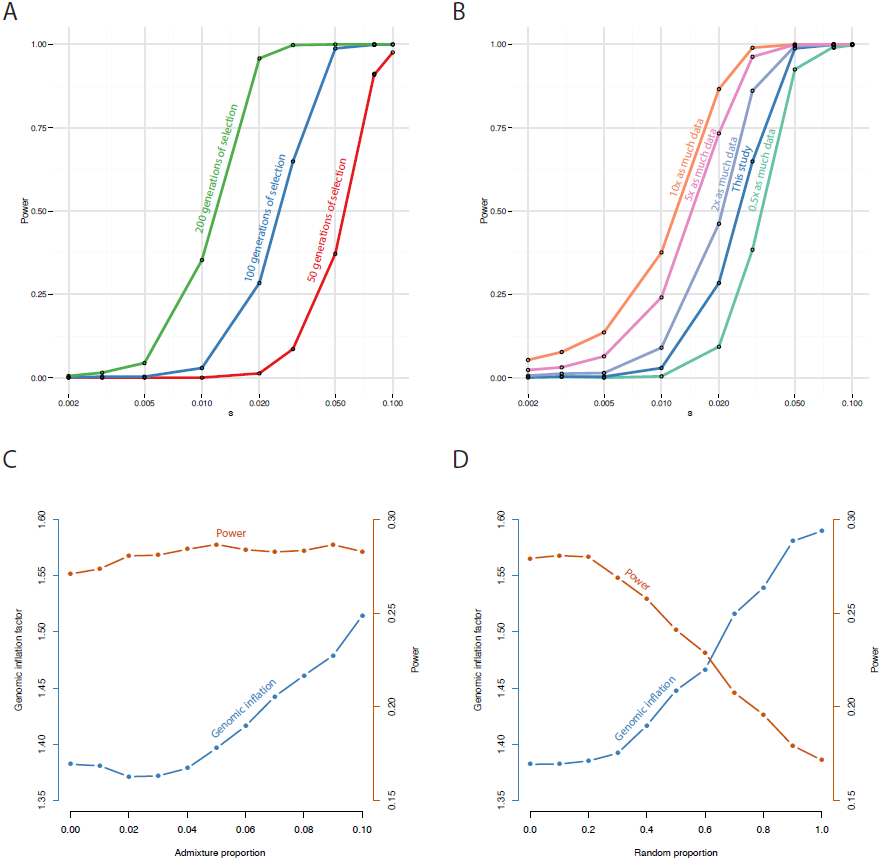
Estimated power of the selection scan. **A:** Estimated power for different selection coefficients for a SNP that is selected in all populations for either 50, 100 or 200 generations. **B:** Effect of increasing sample size, showing estimated power for a SNP selected for 100 generations, with different amounts of data, relative to the main text. **C:** Effect of admixture from Yoruba (YRI) into one of the modern populations, showing the effect on the genomic inflation factor (blue, left axis) and the power to detect selection on a SNP selected for 100 generations with a selection coefficient of 0.02. **D:** Effect of misspecification of the mixture proportions. Here 0 on the x-axis corresponds to the proportions we used, and 1 corresponds to a random mixture matrix.

## Supplementary Information Archaeological context for 83 newly reported ancient samples

### Overview

This note provides archaeological context information for the 83 samples for which genome-wide data is presented for the first time in this study. Descriptions of the context for the remaining ancient samples analyzed in this study have been published in previous publications. Specific references for each sample are given in Supplementary Data Table 1.

### Anatolia Neolithic (n=26 samples)

Since the 1950’s, central and southeastern Anatolia have been the subject of archaeological research focused on the origins of sedentism and food-production. The Marmara region of northwestern Anatolia, the location of the to archaeological sites that are the sources of the Anatolian samples in this study (Menteşe Höyük and Barcın Höyük) was for a long time considered to be a region where early agricultural development had not occurred.

Systematic multidisciplinary research in the eastern Marmara region started at the end of the 1980s under the auspices of the Netherlands Institute for the Near East, Leiden (NINO) and its annex in İstanbul, the Netherlands Institute in Turkey (NIT). The first excavations were at Ilıpınar Höyük near İznik Lake. These were followed by excavations at Menteşe and Barcın, situated near an ancient lake in the basin of Yenişehir. These sites have provided key data on early farming communities, their environmental settings, and their cultural development over more than a thousand years, from 6600 calBCE (Barcın) to 5500 calBCE (Ilıpınar). A fourth important site in the same region, Aktopraklık, is being excavated by the Prehistory Department of Istanbul University.

These Neolithic villages are distinctive for monochrome pottery, a sophisticated bone industry, single room dwellings with mud and timber frame walls, and mud-brick buildings that were either freestanding or side-by-side in rows. Ilıpınar includes a large number of free standing timber wall buildings consisting of posts closely set in ditches and coated with mud, and pitched reed-covered roofs carried by central posts^1^. These buildings are regarded as prototypes for houses in eastern and central Europe after the onset of the European Neolithic.

Another characteristic of the eastern Marmara region is the abundance of graves in or near settlements. Given the few hundred burials uncovered so far, it has been hypothesized that the regional tradition involved interment of a large part of the deceased population in the village ground. This custom contrasts with nearby regions of western Anatolia and Turkish and Bulgarian Thrace, where inhumations are scarce^2^. In central Anatolia, for example at Çatal Höyük^3^, burial frequencies are similar to the Marmara region. There are also other links between the two regions in terms of ceramic assemblages and other material culture elements.

With a few exceptions, the funeral practices of the eastern Marmara region were as follows. Graves were spread across the courtyards or concentrated in one spot near the village^4^. The dead were buried in single graves and lay in contracted positions on their left or right side with varying body orientations. The articulated skeletal parts point to primary interments. Another feature of the burials is the presence of wooden remains in many grave pit bottoms, showing that the deceased had been deposited on wooden boards^5^. Grave goods often accompanied the dead, as there were pottery vessels, necklaces, pendants, stone tools, and also animal parts (horns, scapulae).

#### Mentese Höyük in Anatolia (n=5 samples)

Several archaeological excavations were carried out at Menteşe between 1996 and 2000 with the purpose of comparing the extensive chronological and socio-economic data from Ilıpınar with that from Menteşe. Menteşe was a small farming community in the plain of Yenişehir with a millennium-long occupation history and an economy based on plant cultivation and animal husbandry. Houses were built with light mud and timber-frame walls of the wattle-and-daub type. The material culture includes monochrome pottery and falls within the tradition of the Marmara region loosely defined as the Fikirtepe culture^6^. Eleven radiocarbon dates running from 6400 to 5600 calBCE corroborate the stratigraphy of the mound.

The total number of individuals excavated at Menteşe is 20: 11 adults and 9 infants and children^7^. The following 5 individuals produced genome-wide data:

##### • I0724 / UP

This individual is estimated to be 10-14, years old, with a stature of 157 cm. Osteological analysis indicates a male, and we confirm this genetically. He was found close to an adult woman who was older than 40. He was lying on his left side in a S-N orientation with his back turned to the woman.

##### • I0725 / UA SSK15 (filtered out of main analyses)

This individual is estimated to be 23-34 years old, with a stature of 155 cm. Osteological analysis indicates a female, which we confirm genetically. She had two traumas: a healed impression fracture of the vault of her skull, and a split and fused distal phalanx of one of her thumbs. She was buried on her right side in a SW-NE orientation, and wooden remains at the bottom of her grave indicated she was buried on a wooden plank. Two aspects of her burial were unusual and suggested the possibility of a later burial: her grave yielded some broken pottery of a type that is unusual for the Menteşe archaeological assemblage, and the truncated burial shaft was dug more than one meter deep from an unknown surface. Population genetic analysis of data from this individual indicates that she is an outlier compared to other Anatolian Neolithic samples. Combined with the other unusual features of her burial, this led us to filter her data out of our main analyses.

##### • I0727/UA JK 16

This individual is estimated to be 34-40 years old, with a stature of 168 cm. Osteological analysis indicates a male, which we confirm genetically. He probably suffered from diffuse idiopathic skeletal hyperostosis (DISH) and osteoarthritis, and his left forearm showed a healed fracture. His highly flexed skeleton was buried its right side in a SW-NE orientation. A broken pot that was of a style that does not fit well for the archaeological assemblage of Menteşe was found near his face. Despite this feature, we did not filter out his data from our main analyses because unlike I0725 / UA SSK15, he is not a population genetic outlier.

##### • I0726 / UF

This individual is estimated to be 23-40 years old. Osteological analysis indicates a male, but the genetic data contradicts this and suggests a female. The individual, who suffered from osteoarthritis, was buried with pottery vessels as indicated by sherds near the head and feet, and was lying on its right side in a W-E orientation.

##### • I0723 / UH

This individual is estimated to be 43-48 years old, with a stature of 166 cm. Osteological analysis indicated a male, which we confirm genetically. A bone tool found near his waist suggests that he wore a belt when he was buried. Like I0726 / UF, he suffered from osteoarthritis. His skeleton was lying on its left side in a S-N orientation.

#### Barcin Höyük in Anatolia (n=21 samples)

Barcın Höyük is located in the Yenişehir Plain in Northwest Turkey, originally on a small natural elevation at the edge of a retreating lake^8^. Excavations have demonstrated continuous occupation between around 6600 calBCE and 6000 calBCE^9^, producing about 4 meters of stratified Neolithic settlement deposits. From the start of habitation, animal husbandry and crop cultivation were the mainstay of this subsistence economy. The deepest levels at Barcın Höyük represent the oldest known farming community in northwestern Anatolia.

It is important to place the site of Barcın within the context of a broader understanding of the Neolithization of western and northwestern Anatolia. Ongoing excavations in both regions now indicate a rather sudden appearance of farming villages around 6700-6600 calBCE^10^. This development breached a boundary that had remained in place for more than 1,500 years between agricultural landscapes in southeastern and central Anatolia to the east and uninhabited or lightly inhabited forager landscapes to the west^11^.

Excavations at Barcın between 2007 and 2014 yielded large assemblages of human remains, mostly from primary and single inhumation burials within the settlement^2^. Infants, juveniles and adults are all represented. To date, remains of 115 individuals have been excavated (47 adults, 68 non-adults). More than half of the burials belong to infants. DNA was successfully extracted from 21 samples (15 infants, 1 child, 5 adults). Fourteen of the 21 the skeletons were osteologically analyzed. The age determination of the infants that were not examined was estimated by observing bone dimensions when the samples were collected.

The sample numbers and descriptions are given below:

##### • I0707 / L11-213

This infant is estimated to be about 3 months old based on long bone size and 4-8 months old based on dentition. The sample is genetically determined to be female.

##### • I0708 / L11-439

This is a badly disturbed burial that is both osteologically and genetically male.

##### • I0709 / M13-170

This is an older infant estimated from bone size. The sample is genetically male.

##### • I0736 / L11-216

This is estimated to be a neonate (from the bone size). The sample is genetically female. It is also genetically a first degree relative of I0854 / L11-215.

##### • I0744 / M10-275

This is estimated to be a neonate (from the bone size). The sample is genetically male.

##### • I0745 / M11-363

This is estimated to be a neonate (from the bone size). The sample is genetically male.

##### • I0746 / L11-322

This infant is estimated to be about 3 months old based on long bone size and 4-8 months old based on dentition. The sample is genetically male.

##### • I0854 / L11-215 (filtered out of main analyses)

This is estimated to be a neonate (from the bone size). The sample is genetically female. The sample is also genetically a first degree relative of I0736 / L11-216. We use this sample’s first degree relative to represent this family because it is has more genetic data.

##### • I1096 / M10-76

This infant is estimated to be 1-2 years old. The sample is genetically male.

##### • I1097 / M10-271

This is estimated to be a neonate (from bone size). The sample is genetically male.

##### • I1098 / M10-352

This is estimated to be a neonate (from bone size). The sample is genetically female.

##### • I1099/ L11-S-488

This is estimated to be a neonate (from bone size). The sample is genetically male.

##### • I1100 / M11-351

This is estimated to be a neonate (from bone size). The sample is genetically female.

##### • I1101 / M11-352a

This is estimated to be a neonate (from bone size). The sample is genetically male.

##### • I1102 / M11-354

This is estimated to be a neonate (from bone size). The sample is genetically male.

##### • I1103 / M11-S-350

This is estimated to be a neonate (from bone size). The sample is genetically male.

##### • I1579 / M13-72

This individual is osteologically determined to be a 35-45 year old female, and is also genetically female. She was buried on her left side in a highly flexed position.

##### • I1580 / L12-393

This individual, genetically female, was not fully examined. A flint artefact was in the grave.

##### • I1581/ L12-502

This adult, genetically female, was not examined.

##### • I1583/L14-200

This child, age about 6-10 years, is genetically male.

##### • I1585 / M11-59

This badly disturbed burial of an individual from middle to older age did not have an osteologically determined sex. The sample is genetically female.

### Iberia Chalcolithic (n=14 samples)

#### El Mirador Cave in Iberia (n=14 samples)

All the Iberia Chalcolithic samples are from El Mirador Cave, which overlooks the southernmost flank side of Sierra de Atapuerca (Burgos, Spain), at an altitude of 1,033 meters above sea level. The mouth of this karst cavity is now approximately 23 meters wide and 4 meters high, penetrating some 15 meters inwards. Initial archaeological work was carried out in the 1970s. In 1999, the fieldwork was resumed, and it is still ongoing^12^.

**Figure 1:**
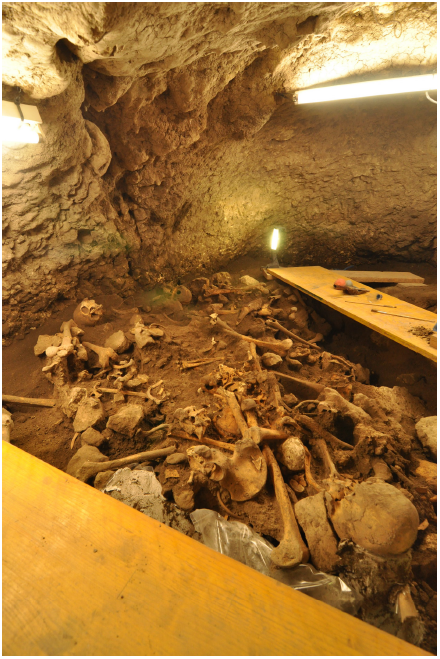
Picture of the collective burial of El Mirador cave, showing the Chalcolithic skeletons analyzed (2014 excavation season).

a. A single burial from the Middle Bronze Age dated to 1,740 calBCE (Beta-296226)
b. Six individuals from the Early Bronze Age dated to between 2,030 and 2,430 calBCE (Beta-153366, 182041 and 182042) that correspond to human remains that were cannibalized and then abandoned^13^.
c. A minimum of 23 individuals from a collective burial from the Chalcolithic period, excavated in a small natural cavity located in the NE corner of the cave (Fig. 1). The remains are associated with smooth hemispherical bowls, fractured deer antlers, and river shell valves. Two bones have been radiocarbon dated, yielding similar dates: 2,880 calBCE (Beta-296227), and 2,630 calBCE (Beta-296225). Based on the union of the calibrated confidence intervals, we estimate the burials to be between 2,570 and 2,900 years calibrated BCE. Mitochondrial DNA of some specimens was published in 2014^14^.

The following 14 samples successfully produced genome-wide data:

##### • I0581 / MIR5 and MIR6

This sample is genetically male. Initially these samples were thought to be from two different individuals but the genetic analysis indicates they are the same individual so we merged them.

##### • I1274 / MIR11 (filtered out of main analyses)

This sample is genetically male. Genetic analysis indicates that this individual is a first degree relative (parent-child or sibling) of I1277 / MIR14. We filtered this sample from our main analyses because its first degree relative has more genetic data.

##### • I1277 / MIR14

This sample is genetically male. Genetic analysis indicates that this individual is a first degree relative (parent-child or sibling) of I1274 / MIR11.

##### • I1302 / MIR24 (filtered out of main analyses)

This sample is genetically male. Genetic analysis indicates that this individual is a first degree relative (parent-child or sibling) of I1314 / MIR26. We filtered this sample from our main analyses because its first degree relative has more genetic data.

##### • I1314 / MIR26

This sample is genetically male. Genetic analysis indicates that this individual is a first degree relative (parent-child or sibling) of I1302 / MIR24.

##### • I1271 / MIR1

This sample is genetically female.

##### • I1272 / MIR2

This sample is genetically female.

##### • I1276 / MIR13

This sample is genetically female.

##### • I1280 / MIR17

This sample is genetically female.

##### • I1281 / MIR18

This sample is genetically female.

##### • I1282 / MIR19

This sample is genetically male.

##### • I1284 / MIR21

This sample is genetically male.

##### • I1300 / MIR22

This sample is genetically female.

##### • I1303 / MIR25

This sample is genetically male.

### Neolithic samples from Mittelelbe-Saale, Germany (n=15)

This study includes 15 newly reported samples from the transect-through-time study in Mittelelbe-Saale, Central Germany that featured in previous publications^15,16^. For the reader’s convenience and reasons of consistency, we repeat summary descriptions for some sites followed by the list of samples for which genome-wide data are newly reported in this study.

#### LBK in Germany: Halberstadt-Sonntagsfeld (n=1)

The *Linearbandkeramik* (LBK) Halberstadt-Sonntagsfeld site was discovered during construction of a new development and was excavated between 1999 and 2002, uncovering 1324 archaeological features across an area of 9947m^2^. The majority of the finds could be attributed to the LBK, with contours and remnants of seven long houses, several pits and 42 graves^17^. The remaining finds could be attributed to the Middle Neolithic Bernburg culture as well as the Unetice and Urnfield cultures of the Bronze Age. The site is a classical example of LBK settlement burials, where the majority of graves were grouped around five long houses, while one group of six graves was located inside the central yard area of the settlement. We add one 40-60 year-old individual clearly associated with an LBK house.

##### • I1550 / HAL19

This is grave 22, feature 666. The sample is osteologically and genetically female.

#### LBK in Germany: Karsdorf (n=1)

The site of Karsdorf is located in the valley of Unstrut, Burgenlandkreis, Saxony-Anhalt, Germany. The slope on which Karsdorf is located is characterized by alluvial loess. The place itself was settled intensively since the earliest phase of the LBK in the region. The settlement area is at least 50 acres in size and nearly 30 houses have been excavated. So-called ‘settlement burials’ were regularly found in pits in the center of the settlement area. We add one individual.

##### • I0797 / KAR16

Feature 611. This sample is genetically male.

#### Salzmuende culture in Germany: Salzmünde-Schiebzig (n=1)

The spur on which the site of Salzmünde (7 km south of Halle/Saale, Saxony-Anhalt) is located has been inhabited intermittently since the Paleolithic. Human occupation intensified during the Neolithic, with archaeological evidence for settlements and burials. Around 3300 BCE people of the eponymous Middle Neolithic Salzmünde culture built a causewayed enclosure (earthwork) with double trenches. Ground plans of houses and several clay pits indicate permanent settlements at the site. Excavation between 2005 and 2008 found human remains of 141 individuals, which reveal a number of complex burial rites: remains were found both inside and outside the enclosure, sometimes underneath layers of potsherds, and sometimes as isolated skeletal elements (mostly skulls) in trenches and clay pits. We report one new individual from this culture.

##### • I0551 / SALZ3b

Feature 6582. This is a 2.5-4 year old child from an unusual multiple grave of nine individuals that was found in a 1-1.5m deep, circular pit of 1.3m diameter underneath a layer of 8000 sherds of amphorae and other pottery. The sample is genetically male.

#### Corded Ware in Germany: Esperstedt (n=9)

The site of Esperstedt forms part of large-scale excavations initiated in 2005 in the context of major infrastructural roadworks in Saxony-Anhalt, Germany to build motorway A38. Individuals from Esperstedt reference site 4 could be unambiguously assigned to the Corded Ware culture, both by accompanying pottery and by characteristic orientation of the burials^18^. Males were usually buried in a right-hand side flexed position with head to the west and facing south, while females were buried on their left-hand side with their head to the east. We added nine new individuals from this site:

##### • I1532 / ESP8

Feature 4182. This is genetically male.

##### • I1534 / ESP14

Feature 6141. This is genetically male.

##### • I1536 / ESP17

Feature 4098. This is genetically male.

##### • I1538 / ESP20

Feature 2200. This is genetically male.

##### • I1539 / ESP25

Feature 4179. This is genetically female.

##### • I1540 / ESP28

Feature 2152. This is genetically male.

##### • I1541 / ESP32

Feature 4290. This is genetically male.

##### • I1542 / ESP33

Feature 2101. This is genetically male.

##### • I1544 / ESP36

Feature 6232. This is genetically male.

#### Bell Beaker in Germany: Benzingerode-Heimburg (n=2)

The archaeological site was excavated 2000-2005 in the context of major infrastructural roadworks in Saxony-Anhalt, Germany, to build motorway B6n. Numerous Late Neolithic finds across the 2.5km long and 30-100m wide section V between the villages Benzingerode and Heimburg indicate settlement activity during the Bell Beaker and Early Bronze Age Unetice periods^19^. We include two new individuals attributed to the Bell Beaker period:

##### • I1546 / BZH2

Grave 8, feature 5079. This individual is genetically female. The body was oriented in right-hand-side flexed position in SO-NW orientation with the head in the southwest facing north. Based on this it was tentatively classified as Bell Beaker by excavator Tanja Autze^20^.

##### • I1549 / BZH15

Grave 1, feature 1289. This individual is osteologically and genetically female, and was classified as Bell Beaker x by excavator Tanja Autze^1^. The body of the 50-65 year-old was oriented in right-hand-side flexed position with the head in the east facing northeast.

#### Bell Beaker in Germany: Quedlinburg VII (n=1)

The site of Quedlinburg, Harzkreis, is situated in the fertile foothills of the northern Harz, a region characterized by rich loess soils. A group of six graves was discovered at the Quedlinburg reference site VII in Saxony-Anhalt and has been attributed to the Bell Beaker culture based on the form and orientation of the burials^21^. We included one new individual:

##### • I0805 / QLB26

Feature 19614. This 35-45 year-old individual is osteologically and genetically male. The body was buried in NO-SW orientation with the head in the north facing east. Grave goods are scarce and include three silex arrowheads, a few potsherds, and animal bones. A notable observation from the physical anthropological examination is traits at the acetabulum and the femur head suggesting that the individual frequently rode horses.

### Prehistoric Samples from Russia, Eneolithic to Bronze Age (n=27)

This study contains 26 additional samples from Russia obtained during the Samara Valley Project, and 1 from Karelia, in addition to those previously published in ref. 16. For the reader’s convenience we repeat summary descriptions for some sites that contained both previously published graves and graves that are published here for the first time.

#### Hunter-gatherer sample: Karelia Russia (n=1)

In this study we added another individual from the ~5500 BCE Mesolithic site Yuzhnyy Oleni Ostrov (an island in Lake Onega) in Karelia, Western Russia, to the one reported in ref. 16. Mitochondrial data from seven other individuals from the same site have been described^22^.

##### • I0221 / UZ0040

MAE RAS collection number 5773-40, grave number 39/1. This is genetically male.

#### Khvalynsk Eneolithic in the Volga steppes: Saratovo, Russia (n=3)

Three individuals described here were among 39 excavated in 1987-88 at the Eneolithic cemetery of Khvalynsk II, Saratov oblast, Russia, on the west bank of the Volga River, 6 km north of the village of Alekseevka. Khvalynsk I and II are two parts of the same cemetery, excavated in 1977-79 (Khvalynsk I) and 1987-88 (Khvalynsk II).^23^ The two excavations revealed 197 graves, about 10x larger than other cemeteries of this period in the Volga-Ural steppes, dated by radiocarbon to 5200-4000 BCE (95.4% confidence). Bones of domesticated cattle and sheep-goat, and horses of uncertain status, were included in 28 human graves and in 10 sacrificial deposits. The 367 copper artifacts in the graves, mostly beads and rings, are the oldest copper objects in the Volga-Ural steppes, and trace elements and manufacturing methods in a few objects suggest trade with southeastern Europe. Together with high ^15^N in the human bones from Khvalynsk, which might have caused a reservoir effect making ^14^C dates too old, the circulation of so much copper, which increased in SE Europe after 4700 BCE, suggests that a date after 4700 BCE would be reasonable for many graves at Khvalynsk. Copper was found in 13 adult male graves, 8 adult female graves, and 4 sub-adult graves. The unusually large cemetery at Khvalynsk contained southern Europeoid and northern Europeoid cranio-facial types, consistent with the possibility that people from the northern and southern steppes mingled and were buried here.

##### • 10122 / SVP35 (grave 12)

Male (confirmed genetically), age 20-30, positioned on his back with raised knees, with 293 copper artifacts, mostly beads, amounting to 80% of the copper objects in the combined cemeteries of Khvalynsk I and II. Probably a high-status individual, his Y-chromosome haplotype, R1b1, also characterized the high-status individuals buried under kurgans in later Yamnaya graves in this region, so he could be regarded as a founder of an elite group of patrilineally related families. His MtDNA haplotype H2a1 is unique in the Samara series.

##### • 10433 / SVP46 (grave 1)

Male (confirmed genetically), age 30-35, positioned on his back with raised knees, with a copper ring and a copper bead. His R1a1 haplotype shows that this haplotype was present in the region, although it is not represented later in high-status Yamnaya graves. His U5a1i MtDNA haplotype is part of a U5a1 group well documented in the Samara series.

##### • 10434 / SVP47 (grave 17)

Male (confirmed genetically), age 45-55, positioned contracted on his side, with 4 pathological wounds on his skull, one of which probably was fatal. No grave gifts or animal sacrifices accompanied the burial. His Q1a Y-chromosome haplotype is unique in the Samara steppe series, but his U4a2 or U4d MtDNA haplotype are not unusual.

#### Middle Bronze Age Poltavka culture, Samara, Russia (n=5)

Five Middle Bronze Age (MBA) individuals described here were excavated by Samara archaeologists at five different MBA kurgan cemeteries of the Poltavka culture, all located in the Samara oblast. The Poltavka culture evolved from the Yamnaya culture beginning about 2900-2800 BCE and lasted until about 2200-2100 BCE. Four of the five individuals are directly dated with ^14^C dates that fall between 2925-2484 BCE. Poltavka graves exhibited new types of pottery vessels and some small innovations in grave shape and construction compared with the earlier Yamnaya, but the Poltavka economy was a continuation of the mobile pastoral economy introduced during the Early Bronze Age Yamnaya period beginning about 3300 BCE. MBA Poltavka settlements excavated during the Samara Valley Project were ephemeral seasonal camps that yielded <1 artifact per 2m^2^, a signal of high mobility.

##### Kutuluk III, kurgan 1, grave 2

###### • 10126 / SVP51

Kutuluk III is a Poltavka kurgan cemetery dated to the MBA located on the Kinel’ River, a tributary of the Samara, near two earlier EBA Yamnaya kurgan cemeteries. Kurgan 1, grave 2 contained a male (confirmed genetically), age 25-35, feet stained with red ochre, ovicaprid bones associated in the grave. Y-chromosome haplotype R1b1a2a2, MtDNA type H6a2.

##### Grachevka II, kurgan 1, grave 1

###### • 10371 / SVP11

Grachevka II is a Poltavka kurgan cemetery dated to the MBA located on the Sok River, north of the Samara River in Samara oblast, within a regional cluster of Bronze Age kurgan cemeteries. Kurgan 1, grave 1 contained a male (confirmed genetically), age not recorded. Y-chromosome haplotype R1b1a2, MtDNA type U2d2.

##### Nikolaevka III, kurgan 5, grave 1

###### • 10374 / SVP16

Nikolaevka is a cemetery of five kurgans dated to the MBA located south of the estuary of the Samara River near its junction with the Volga. Kurgan 5, grave 1 contained a male (confirmed genetically) age 35-45. Y-chromosome haplotype R1b1a2a, MtDNA type H13a1a.

##### Lopatino II, grave kurgan 1, grave 1

###### • 10440 / SVP53

Lopatino II is a cemetery of kurgans dated to both the EBA and MBA located on the Sok River in Samara oblast, within a regional cluster of Bronze Age kurgan cemeteries. Kurgan 1, grave 1 contained a male (confirmed genetically) age 25-35 with red ochre staining on his distal tibia. Y-chromosome haplotype R1b1a2a2, MtDNA type I3a.

##### Potapovka I, kurgan 5, grave 6

###### • 10432 / SVP42

Potapovka I is an important cemetery of the late MBA or MBA2 Sintashta-culture era, but this grave shows that an older MBA Poltavka cemetery was located in the same place on the Sok River, in the transitional forest-steppe zone of Samara oblast. Grave 6 was dated 2925-2536 calBCE (4180 ± 84 BP: AA12569), centuries older than the MBA2 grave pit that cut through it, removing about 60% of the Grave 6 skeleton. Grave 6 was that of a male (confirmed genetically) age 35-45 years, his foot bones stained with red ochre, buried with the lower leg bones of a sheep or goat. His Y-chromosome haplotype was R1a1a1b2a, the only R1a male in the EBA-MBA series and a signal of an emerging broader set of more varied male haplogroups that would become better defined in the MBA2 and LBA. His MtDNA haplotype was U5a1c. Another Poltavka grave under kurgan 3 was cut through in the same way, so Potapovka 1 seems to have been established in the MBA2 directly on top of an older Poltavka cemetery.

#### Late Middle Bronze Age (MBA2) Potapovka culture, Samara, Russia (n=3)

Three individuals of the MBA2 period, 2200-1800 calBCE, were excavated from two MBA2 kurgan cemeteries of the Potapovka culture near the village of Utyevka in the Samara oblast. The Potapovka culture was a western variant of the Sintashta culture, which was centered 400 km to the east and produced the oldest dated chariots, buried in graves with paired horse teams and weapons. In the Samara steppes, cemeteries of the Potapovka type contained material culture very similar to Sintashta in graves with paired horses and cheekpieces, but no spoked wheel remains survived.^24^ Sintashta and Potapovka are argued to have been the parent cultures from which the LBA Andronovo and Srubnaya cultures evolved, and so played a key role in the transition from the mobile EBA/MBA economies to the more settled LBA economy. The MBA2 period witnessed innovations in warfare and weaponry, the beginning of a change to sedentary pastoralism, and the extension of trade contacts from the northern steppes to Central Asian urban centers. The population afforded graves under kurgans expanded to include not just a few adult men, but many women and children as well, perhaps entire elite families.

##### Utyevka VI, kurgan 6, grave2

###### • 10246 / SVP41

The Utyevka VI cemetery was located 0.8 km north-northeast of the village of Utyevka, Samara oblast, south of the Samara River. It contained some of the richest and most unusual graves of the Potapovka culture. Kurgan 6, grave 2 contained six humans: a male-female couple buried facing each other aged 15-17; and four children and infants too old and numerous to be the offspring of the couple—perhaps siblings. Horse sacrifices, shield-shaped studded bone cheekpieces interpreted as chariot-driving gear, weapons (three copper daggers, a flat copper axe, 16 flint projectile points), copper beads and rings, and other objects were found in the grave. Sample 10246 is from the 15-17-year-old male (confirmed genetically). His Y-chromosome haplotype was P1.

##### Utyevka IV, kurgan 4, grave 1

###### • 10418 / SVP24

The Utyevka IV cemetery was located 1 km SE of the Utyevka VI cemetery. Kurgan 4, grave 1 contained a mature female (confirmed genetically) age 45+, buried with the bones of an ovicaprid. Her MtDNA haplotype was T1a1.

##### Utyevka VI, kurgan 7, grave 1

###### • 10419 / SVP27

Kurgan 7, grave 1 at Utyevka VI contained a mature male (confirmed genetically) age 45+. He was buried with a Potapovka-style ceramic pot and his grave was covered by a pair of sacrificed horse skulls and hooves probably originally attached to their hides. His Y-haplotype was R1a1a1b; MtDNA was U2e1h, found in other ancient steppe graves.

#### Late Bronze Age Srubnaya culture, Samara, Russia (n=14)

Fourteen individuals of the LBA Srubnaya, or Timber-Grave, culture were sampled from 7 kurgan cemeteries in the middle Volga region of Russia. The LBA Srubnaya-Andronovo horizon was a chain of similar settlements, cemeteries, and material culture types associated with a settled form of pastoralism that extended across the Eurasian steppe belt, connecting China with Europe and Iran for the first time. The upper Samara valley south of the Urals was the border between Srubnaya to the west and Andronovo to the east; many settlements in this region contain both styles of pottery. Srubnaya material culture types appeared earliest about 1900 BC in the Samara region, then spread west and south to the Dnieper in Ukraine by 1700 BC. Srubnaya settlements extended north into the forest-steppe zones in the Samara region between 1900-1200 BC. Srubnaya cemeteries included the whole population in normal proportions by age and sex. Unlike the cemeteries of the EBA-MBA and MBA2, the cemeteries of the LBA provide a demographic sample with data in all age and sex classes. Chariots were used, and large copper mines operated in and around Samara oblast, although most Srubnaya settlements in the region were family-sized homesteads of 3-5 structures. The Spiridonovka II cemetery was the richest Srubnaya cemetery in the Samara valley, probably associated with a local chiefly family.

##### Novosel’ki, kurgan 6, grave 1

###### • 10232 / SVP12

Novosel’ki was a Srubnaya cemetery containing 11 kurgans and 76 Srubnaya graves, located in Tatarstan oblast near the forest-steppe/forest zone ecotone, an early example of a Srubnaya community located in the forest-steppe zone, well north of the steppes. Kurgan 6, grave 4 contained a young male (confirmed genetically) age 17-25, with a normal steppe Y-chromosome haplotype of R1a1a1b2, MtDNA type U5a1f2.

##### Rozhdestvenno I, kurgan 5, grave Z

###### • 10234 / SVP25

Rozhdestveno was another forest-steppe zone Srubnaya cemetery located west of the Volga. Kurgan 5 grave 7 contained an adult female (confirmed genetically) age 25-35, MtDNA type K1b2a, previously unusual in the steppes.

##### Rozhdestvenno I, kurgan 4, grave 4, skeleton 2

###### • 10235 / SVP26

Rozhdestveno I, kurgan 4, grave 4, skeleton 2 was that of an adolescent female (confirmed genetically) age 15-17. Her MtDNA haplotype was I1a1, previously unusual is the steppes.

##### Barinovka I, kurgan 2, grave 24

###### • 10422 / SVP30

Adult female (confirmed genetically) age 40-45, MtDNA type T1a1.

##### Barinovka I, kurgan 2, grave 17

###### • 10423 / SVP31

Adult male (confirmed genetically) age 40-50 with Y-chromosome type R1a1a1b2; MtDNA type was J2b1a2a.

##### Uvarovka I, kurgan 2, grave 1

###### • 10424 / SVP32

Adult male (confirmed genetically) age 40-50 with Y-chromosome type R1a1a1b2; MtDNA type was T2b4.

##### Spiridonovka IV,

Spiridonovka IV was one of four Srubnaya kurgan cemeteries located on the southern bank of the Samara River, north and south of the village of Spiridonovka. Cemetery IV was located southeast of the village and contained 9 kurgans, two of which were excavated by Samara archaeologists in 1996, uncovering 24 Srubnaya individuals in kurgan 1 and 15 in kurgan 2, including 7 adult males, 5 adult females, 11 sub-adults, and 1 undetermined. About half the graves contained a pottery vessel and a few had beads or a bracelet of copper or bronze.

##### Spiridonovka IV, kurgan 1, grave 15

###### • 10354 / SVP1

Adult female (confirmed genetically) age 35-45 with MtDNA type U5a1.

##### Spiridonovka IV, kurgan 2, grave 1

###### • 10358 / SVP6

Adult female (confirmed genetically) age 35-35 with MtDNA haplotype H6a1a.

##### Spiridonovka IV, kurgan 1, grave 6

###### • 10359 / SVP7

Adult female (confirmed genetically) age 25-35, MtDNA haplotype U5a2a1.

##### Spiridonovka IV, kurgan 1, grave 11

###### • 10360 / SVP8

Male (confirmed genetically) of undetermined age; a secondary burial of bones exposed and only partially collected before burial. His Y-chromosome haplotype was R1a1; his MtDNA type was U5a1.

##### Spiridonovka IV, kurgan 2, grave 5

###### • 10361 / SVP9

Adolescent male (confirmed genetically) age 15-17, Y-chromosome type R1a1a, MtDNA type H5b.

##### Spiridonovka II

Spiridonovka II was the richest Srubnaya cemetery in the lower Samara River valley, located northwest of the village of Spiridonovka. At least 14 kurgans were counted, and four were excavated (1, 2, 10, 11). Kurgan 1 contained 17 graves and kurgan 2, 37 graves. Grave 1 in kurgan 1 contained a male age 17-25 with weapons—a dagger and a set of 12 lanceolate flint projectile points—and a bone clasp or belt ornament. Grave 2 contained an adolescent age 13-16, probably female, and a juvenile age 6-8 years, decorated with faience beads that were probably imported from outside the steppes, and numerous bronze bracelets, beads, and pendants, including a pair of long, dangling earrings made of leaf-like sets of pendants arranged in three rows. Grave 7 contained a bronze bracelet, ring, and bead, while Grave 13 contained an adult female age 25-35 with 16 tubular bronze beads and eight bronze medallions. In kurgan 2, grave 29 contained a child about 12 years of age buried with a copper-bronze bracelet and a pair of gold-silver pendants gilded using a technique known as depletion gilding, documented elsewhere before the first millennium BC only in the jewelry from the Royal Cemetery of Ur in Iraq. Samples 10421 and 10430 below, buried under different kurgans with distinct groups of graves, were first-degree relatives, and given their shared MtDNA types, they were either mother and son or brother and sister.

##### Spiridonovka II, kurgan 11, grave 12

###### • 10421 / SVP29

Adult female (confirmed genetically) 35-45, MtDNA type H3g. She is a first degree relative of I0430. We exclude her data from population genetic and selection analyses because of the higher coverage of their first degree relative, who we use to represent the family.

##### Spiridonovka II, kurgan 1, grave 1

###### • 10430 / SVP39

Adult male (confirmed genetically) age 17-25 buried with weapons, see above; Y-chromosome type R1a1a1b2a2a; MtDNA type H3g. He is a first degree relative of I0421.

##### Spiridonovka II, kurgan 1, grave 2

###### • 10431 / SVP40

Adolescent age 13-16, female (confirmed genetically), MtDNA type H2b.

## Supplementary Information section 2

### Population interactions between Anatolia, mainland Europe, and the Eurasian steppe

Substantial progress has been made in the study of ancient European population history due to the ability to obtain genome-wide data from ancient skeletons^1–15^. The most recent synthesis of European prehistory 8,000-3,000 years ago^5^ presents the following reconstruction.

Before the advent of farming, Europe was populated by at least three different groups of hunter-gatherers. Western European hunter-gatherers^7^ (WHG) were widely distributed in Luxembourg^7^, Iberia^8,11^, and Hungary^3^. Eastern European hunter-gatherers (EHG) lived in far eastern Europe (Russia), including both the steppe (Samara region) and Karelia^5^. Scandinavian hunter-gatherers (SHG) from Sweden^5,7,14^ can be modeled as a mixture of WHG and EHG^5^ and persisted in Scandinavia until after ~5,000 years ago^14,15^. EHG are distinguished by a greater affinity than either WHG or SHG^5^ to Native Americans and to the ~24,000-year old Mal’ta (MA1) individual from Siberia^10^, making them a proximate source for the “Ancient North Eurasian” ancestry present in subsequent Europeans^5,7,16,17^. All European hunter-gatherers were outside the range of genetic variation in present-day Europeans^5,7,8,11,14,15^, but are genetically closest to present-day northern Europeans.

Both the Tyrolean Iceman^6^ and a Funnel Beaker individual from Sweden^15^ (both ~5,000 years old) were at the southern edge of present-day European genetic variation, resembling living Sardinians, rather than the present day populations of the Alps or Scandinavia. The Stuttgart female^7^ (~7,500 years ago) was an early European farmer (EEF) from the Linearbandkeramik (LBK) culture of central Europe and shared this affinity to Sardinians, documenting a relatively homogeneous population of early farmers since the Early Neolithic. The EEF traced part of their ancestry to the Near East but there is substantial uncertainty about the proportion due to a lack of an appropriate reference population^7^. Genome-wide data of Epicardial individuals from Spain^5^ proved that EEF in both Iberia and central Europe belonged to a common ancestral population, with the implication that they were descended from a common source prior to setting along different paths (Danubian and Mediterranean) into Europe. This finding was further confirmed by the analysis of a Cardial individual from Mediterranean Spain^9^.

During the Middle Neolithic period, European farmers in both central Europe and Iberia manifested a “resurgence” of WHG ancestry relative to their Early Neolithic predecessors^5^. The evidence of resurgence of WHG ancestry could reflect a history in which pockets of populations with substantial hunter-gatherer genetic heritage persisted across Europe long after the advent of farming^18^ and then mixed with local farmers in many locations. Alternatively, the resurgence could reflect large-scale population turnover within Europe, whereby at least one population that had a higher proportion of WHG ancestry than many preceding farmer populations, spread and replaced earlier groups.

Unlike Europeans from the Middle Neolithic period and earlier, present-day Europeans cannot be modeled as two-way mixtures of European hunter-gatherers and EEF in any proportions, but instead have at least three sources of ancestry^7^. The third source is related to the MA1 individual from Siberia. This ancestry was conveyed via the EHG and the later Yamnaya steppe pastoralist populations of ~5,000 years ago^5^. It is estimated that steppe ancestry accounts for ~3/4 of the ancestry of early Corded Ware people of the Late Neolithic period ~4,500 years ago^5^ and a substantial portion of ancestry of other Late Neolithic/Bronze Age groups of central Europe (~4,500-3,000 years ago) and present-day Europeans throughout Europe, especially in the north^5^. By contrast, in the eastern expansion of steppe groups to the Altai, represented by the Afanasievo culture^1^, there seems to have been no admixture with local populations. The later Sintashta and Andronovo populations were descended from the earlier steppe populations, but despite living far to the east of the Yamnaya from Kalmykia^1^ and Samara^5^, they had an affinity to more western populations from central and northern Europe like the Corded Ware and associated cultures^1^.

Here we co-analyze 147 previously unreported samples, with 83 samples reported for the first time here that span six archaeological cultures with no published ancient DNA data, to address open questions about the population history of Europe, the steppe, and Anatolia^19^.

### The Anatolian Neolithic is a likely source of the European Neolithic

Our population sample from the Neolithic of Northwestern Anatolia (Fig. 1, Extended Data Fig. 2) has a clear affinity to early European farmers. However, EEF are shifted slightly towards the direction of the WHG in the PCA (Fig. 1) and share more alleles with them than do the Anatolian Neolithic samples (Extended Data Table 3).

To quantify admixture, we used the method described in supplementary information section 9 of Ref. 5, which fits the model:

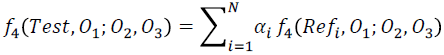

The *Test* population is modeled as an N-way mixture of *Ref_i_* populations in proportions *α_i_*, using *f*_4_-statistics that relate the *Test* and *Ref_i_* populations to a triple of outgroups *O*_1_*; O*_2_, *O*_3_. For m outgroups, there are 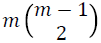 equations of the above form and the proportions *α_i_* are estimated by least squares under the constraint that they sum up to 1 and are in [0, 1]. We use a set of 15 world outgroups^5,7^:

*“World Foci 15” set of outgroups:* Ami, Biaka, Bougainville, Chukchi, Eskimo, Han, Ju_hoan_North, Karitiana, Kharia, Mbuti, Onge, Papuan, She, Ulchi, Yoruba

To enhance this set’s ability to differentiate between reference populations, we can add to it populations that are differentially related to them. For example, when we study admixture in LBK_EN between Anatolia_Neolithic and WHG, we can add EHG as an additional outgroup, as *f*_4_(WHG, Anatolia_Neolithic; EHG, Mbuti)= 0.00823 (Z=22.2), so allele sharing with EHG helps distinguish between WHG and Anatolia_Neolithic as sources of ancestry.

Extended Data Fig. 3C shows that Early Neolithic groups from Europe can be modeled as predominantly of Anatolia_Neolithic ancestry with only 7-11% WHG ancestry. This confirms the visual impression from the PCA (Fig. 1) and ADMIXTURE analysis (Extended Data Fig. 2) of the close relationship between Early Neolithic Europe and Neolithic Anatolia. The relationship is also supported by the low F_ST_=0.005±0.00046 (Supplementary Data Table 2) between LBK_EN and Anatolia_Neolithic and 0.006±0.00054 between Hungary_EN and Anatolia_Neolithic. A direct link between Neolithic Europe and Anatolia is furnished by the fact that around half of the Y-chromosomes of Anatolia_Neolithic belong to haplogroup G2a typical of ancient EEF samples^5,20^, and by the occurrence of mtDNA haplogroup N1a which is typical of EEF^21,22^ in Neolithic Anatolia (Supplementary Data Table 1).

Our results add to the evidence in favor of the hypothesis that the Anatolian Neolithic was the source of the European Neolithic, but we add two notes of caution. First, the fact that our samples are from northwestern Anatolia should not be taken to imply that the Neolithic must have entered Europe from that direction: it simply places a plausible ancestor population at the doorstep of Europe and confirms the hypothesis that the genetic similarity between central European and Iberian farmers could be explained by such a common ancestor. Second, we do not know the geographical extent of people with this ancestry within Anatolia itself and across the Near East. If the population of which the Anatolian Neolithic was a part was sufficiently widespread this ancestry could have entered Europe from a different geographic route. To understand the geographic spread and deeper origin of the Anatolian farmers, it will be necessary to obtain ancient genomes from multiple locations within the Near East.

### Lack of steppe ancestry in the Iberian Chalcolithic

Ancient North Eurasian ancestry is ubiquitous across present-day Europe^5,7^ but was absent in Early and Middle Neolithic Europe^5^, raising the question of when it spread there. When we add Yamnaya_Samara as a third ancestral population there is no improvement in fit for the Iberia_Chalcolithic population (Extended Data Fig. 3B), and this population can be modeled as a mixture of ~77%/23% Anatolia_Neolithic/WHG (Extended Data Fig. 3C). The Iberian Chalcolithic did not have more WHG ancestry than the earlier Middle Neolithic population, as the statistic *f*_4_(Iberia_MN, Iberia_Chalcolithic; WHG, Chimp) is not significantly different from zero (Extended Data Table 3). A recent analysis of a different Iberian Chalcolithic population also suggests that it had more hunter-gatherer ancestry than the earlier European farmers^4^.

The Iberian Chalcolithic population lacks steppe ancestry, but Late Neolithic central and northern Europeans have substantial such ancestry (Extended Data Fig. 3E) suggesting that the spread of ANE/steppe ancestry did not occur simultaneously across Europe. All present-day Europeans have less steppe ancestry than the Corded Ware^5^, suggesting that this ancestry was diluted as the earliest descendants of the steppe migrants admixed with local populations. However, the statistic *f*_4_(Basque, Iberia_Chalcolithic; Yamnaya_Samara,Chimp)=0.00168 is significantly positive (Z=8.1), as is the statistic *f*_4_(Spanish, Iberia_Chalcolithic; Yamnaya_Samara, Chimp)= 0.00092 (Z=4.6). This indicates that steppe ancestry occurs in present-day southwestern European populations, and that even the Basques cannot be considered as mixtures of early farmers and hunter-gatherers without it^4^.

### Transformations of steppe populations

Our paper presents a complete transect of the Samara region beginning with the Samara EHG hunter-gatherer (~5,600BCE)^5^ and ending with the Srubnaya culture (~1,850-1,200 BCE) and a singleton “Scythian” Iron Age individual (~300BCE). In eastern Europe outside the steppe, a new individual from the Karelia region resembles the two previously published EHG individuals^5^ autosomally, but surprisingly belongs to Y-chromosome haplogroup J usually associated with Near Eastern populations (Supplementary Data Table 1).

In a previous study^5^ it was shown that Yamnaya populations from the Samara region were a mixture of the EHG and a population of Near Eastern ancestry for whom present-day Armenians could be a surrogate. The Samara_Eneolithic from Khvalynsk II (~5,200-4,000BCE) predates the Yamnaya by at least 1,000 years but had already begun admixing with this population, although the individuals of this population appear to be heterogeneous (Fig. 1) between EHG and Yamnaya. Taken as a whole, we estimate that they have ~74% EHG and ~26% Armenian related ancestry. The three Eneolithic individuals belong to Y-chromosome haplogroups R1a, R1b, and Q1a; the last of these three is associated with present-day Siberian populations and Native Americans, while R1a and R1b were previously found in the two EHG hunter-gatherers^5^. These results suggest a great degree of continuity with the EHG for the Eneolithic population.

The Yamnaya samples from Samara^5^ and Kalmykia^1^ and the Afanasievo^1^ population from the Altai form a tight cluster (Fig. 1, Extended Data Fig. 2). Our study includes new data from the later Middle Bronze Age population of the Poltavka culture (~2,900-2,200BCE), which cluster with the Yamnaya and Afanasievo. The Poltavka also possess R1b Y-chromosomes (Supplementary Data Table 1), continuing the dominant Y-chromosome type found in the Yamnaya^1,5^. This group of populations have even less EHG ancestry than the Eneolithic population (Extended Data Table 3), estimated to be ~42-52% (Extended Data Fig. 3D).

Admixture into steppe populations continued with the Potapovka culture (~2,500-1,900BCE) and the Srubnaya culture (~1,850-1,200 BCE). Admixture in these later steppe populations is from a different source than in the earlier ones. For the Yamnaya/Afanasievo/Poltavka *Steppe* group, the statistics *f*_4_(EHG, *Steppe;* Armenian, LBK_EN) are negative, and the statistics *f*_4_(*Steppe,* Srubnaya; Armenian, LBK_EN) are positive, suggesting a different source of population change during the EHG→*Steppe* transition and the later Steppe→Srubnaya transition. The *Steppe* group had ancestry related to Armenians and the Srubnaya had an additional source related to European farmers. This is also clear from the PCA where the Srubnaya differ from the *Steppe* group in the direction of European farmers, and in the ADMIXTURE analysis (Extended Data Fig. 2), which shows them to have an EEF/Anatolia Neolithic-related component of ancestry not present in the *Steppe* group. Clearer evidence of this discontinuity is seen when we model different steppe populations as mixtures of Armenians and EHG (Extended Data Fig. 3A), and add Anatolia_Neolithic as a third ancestral population: this has no effect in fit for the earlier populations, but significantly improves fit for the Potapovka, Srubnaya, and eastern Sintashta and Andronovo populations. A discontinuity between earlier and later steppe populations is also suggested by the shift from an R1b Y-chromosome gene pool into an R1a-dominated one in the Srubnaya (Supplementary Data Table 1). We caution that this does not mean that new populations migrated into the steppe as R1a was also detected in Eneolithic Samara and an outlier Poltavka individual (Supplementary Data Table 1); it is possible that R1a males continued to abide in the Samara region but were not included in the rich burials associated with the Yamnaya and Poltavka elites in the intervening period.

It is unclear how the Srubnaya acquired farmer ancestry. One possibility is that contact between early farmer and steppe populations produced populations of mixed ancestry that migrated eastward to the Samara district and further east to form the Sintashta/Andronovo populations. A different possibility suggested in Ref. 1 is that the Corded Ware population of central/northern Europe migrated into the steppe. Our new data document the existence of farmer-admixed steppe populations in the European steppe and provide a plausible source for the more eastward migrations of such populations. Further evidence for a connection between the Srubnaya and populations of central/south Asia—which is absent in ancient central Europeans including people of the Corded Ware culture and is nearly absent in present-day Europeans^23^—is provided by the occurrence in four Srubnaya and one Poltavka males of haplogroup R1a-Z93 which is common in present-day central/south Asians and Bronze Age people from the Altai^24^ (Supplementary Data Table 1). This represents a direct link between the European steppe and central/south Asia, an intriguing observation that may be related to the spread of Indo-European languages in that direction.

Our results on European steppe populations highlight the complexity of their interactions with surrounding populations. At the earliest period, gene flow from a population related to Armenians, presumably from the south, diluted their EHG ancestry and created the mix of ancestry of the Yamnaya/Afanasievo/Poltavka group. Later, gene flow from a population related to Anatolian and European farmers further diluted their EHG ancestry to create the Srubnaya/Sintashta/Andronovo group. This latter group resembled Late Neolithic/Bronze Age populations from mainland Europe (Fig. 1), and like them could be derived from both the farmers of Europe and Anatolia and the earlier steppe populations (Extended Data Fig. 3E). The population history of mainland Europe and the steppe could be summarized as: (i) the dilution of hunter-gatherer ancestry by migrations from different parts of the Near East, (ii) the formation of populations of mixed hunter-gatherer/Near Eastern ancestry (Middle Neolithic/Chalcolithic in mainland Europe, and Eneolithic/Yamnaya/Poltavka on the steppe), and (iii) the migration of steppe populations into Europe during the Late Neolithic (~2,500BCE) and of farmer populations into the steppe, leading to the formation of an array of populations of mixed ancestry encompassing both mainland Europe and the Eurasian steppe. Future research must elucidate how present-day populations emerged from the populations of the Bronze Age, and how populations from mainland Europe, the Eurasian steppe, and Anatolia influenced and were influenced by those from the Near East and Central/South Asia.

## References

1 Grossman, S. R. et al. Identifying recent adaptations in large-scale genomic data. Cell 152, 703–713 (2013).

2 Wilde, S. et al. Direct evidence for positive selection of skin, hair, and eye pigmentation in Europeans during the last 5,000 y. Proc. Natl. Acad. Sci. U. S. A. 111, 4832–4837 (2014).

3 Gamba, C. et al. Genome flux and stasis in a five millennium transect of European prehistory. Nat Commun 5, 5257 (2014).

4 Lazaridis, I. et al. Ancient human genomes suggest three ancestral populations for present-day Europeans. Nature 513, 409–413 (2014).

5 Allentoft, M. E. et al. Population genomics of Bronze Age Eurasia. Nature 522, 167–172 (2015).

6 Keller, A. et al. New insights into the Tyrolean Iceman’s origin and phenotype as inferred by whole-genome sequencing. Nat Commun 3, 698 (2012).

7 Haak, W. et al. Massive migration from the steppe was a source for Indo-European languages in Europe. Nature 522, 207–211 (2015).

8 Olalde, I. et al. Derived immune and ancestral pigmentation alleles in a 7,000-year-old Mesolithic European. Nature 507, 225–228 (2014).

9 Pinhasi, R. et al. Optimal Ancient DNA Yields from the Inner Ear Part of the Human Petrous Bone. PLoS One 10, e0129102 (2015).

10 Alexander, D. H., Novembre, J. & Lange, K. Fast model-based estimation of ancestry in unrelated individuals. Genome Res. 19, 1655–1664 (2009).

11 Patterson, N. et al. Ancient admixture in human history. Genetics 192, 1065–1093 (2012).

12 Underhill, P. A. et al. The phylogenetic and geographic structure of Y-chromosome haplogroup R1a. Eur. J. Hum. Genet. 23, 124–131 (2015).

13 The 1000 Genomes Project Consortium. A global reference for human genetic variation. Nature **In press** (2015).

14 Enattah, N. S. et al. Identification of a variant associated with adult-type hypolactasia. Nat. Genet. 30, 233–237 (2002).

15 Burger, J., Kirchner, M., Bramanti, B., Haak, W. & Thomas, M. G. Absence of the lactase-persistence-associated allele in early Neolithic Europeans. Proc. Natl. Acad. Sci. U. S. A. 104, 3736–3741 (2007).

16 Teslovich, T. M. et al. Biological, clinical and population relevance of 95 loci for blood lipids. Nature 466, 707–713 (2010).

17 Wang, T. J. et al. Common genetic determinants of vitamin D insufficiency: a genome-wide association study. Lancet 376, 180–188 (2010).

18 Price, A. L. et al. The impact of divergence time on the nature of population structure: an example from Iceland. PLoS Genet. 5, e1000505 (2009).

19 Huff, C. D. et al. Crohn’s disease and genetic hitchhiking at IBD5. Mol. Biol. Evol. 29, 101–111 (2012).

20 Hunt, K. A. et al. Newly identified genetic risk variants for celiac disease related to the immune response. Nat. Genet. 40, 395–402 (2008).

21 Jostins, L. et al. Host-microbe interactions have shaped the genetic architecture of inflammatory bowel disease. Nature 491, 119–124 (2012).

22 Beleza, S. et al. Genetic architecture of skin and eye color in an African-European admixed population. PLoS Genet. 9, e1003372 (2013).

23 Barreiro, L. B. et al. Evolutionary dynamics of human Toll-like receptors and their different contributions to host defense. PLoS Genet. 5, e1000562 (2009).

24 Kamberov, Y. G. et al. Modeling recent human evolution in mice by expression of a selected EDAR variant. Cell 152, 691–702 (2013).

25 Turchin, M. C. et al. Evidence of widespread selection on standing variation in Europe at height-associated SNPs. Nat. Genet. 44, 1015–1019 (2012).

26 Berg, J. J. & Coop, G. A population genetic signal of polygenic adaptation. PLoS Genet. 10, e1004412 (2014).

27 Lango Allen, H. et al. Hundreds of variants clustered in genomic loci and biological pathways affect human height. Nature 467, 832–838 (2010).

28 Speliotes, E. K. et al. Association analyses of 249,796 individuals reveal 18 new loci associated with body mass index. Nat. Genet. 42, 937–948 (2010).

29 Heid, I. M. et al. Meta-analysis identifies 13 new loci associated with waist-hip ratio and reveals sexual dimorphism in the genetic basis of fat distribution. Nat. Genet. 42, 949–960 (2010).

30 Morris, A. P. et al. Large-scale association analysis provides insights into the genetic architecture and pathophysiology of type 2 diabetes. Nat. Genet. 44, 981–990 (2012).

31 Briggs, A. W. et al. Removal of deaminated cytosines and detection of in vivo methylation in ancient DNA. Nucleic Acids Res. 38, e87 (2010).

32 Fu, Q. et al. DNA analysis of an early modern human from Tianyuan Cave, China. Proc. Natl. Acad. Sci. U. S. A. 110, 2223–2227 (2013).

33 Fu, Q. et al. An early modern human from Romania with a recent Neanderthal ancestor. Nature (2015).

34 Korneliussen, T. S., Albrechtsen, A. & Nielsen, R. ANGSD: Analysis of Next Generation Sequencing Data. BMC Bioinformatics 15, 356 (2014).

35 International HapMap Consortium. A second generation human haplotype map of over 3.1 million SNPs. Nature 449, 851–861 (2007).

36 Lappalainen, T. et al. Transcriptome and genome sequencing uncovers functional variation in humans. Nature 501, 506–511 (2013).

37 Li, J. Z. et al. Worldwide human relationships inferred from genome-wide patterns of variation. Science 319, 1100–1104 (2008).

38 Loh, P. R. et al. Inferring admixture histories of human populations using linkage disequilibrium. Genetics 193, 1233–1254 (2013).

39 Chang, C. C. et al. Second-generation PLINK: rising to the challenge of larger and richer datasets. GigaScience 4 (2015).

40 Skoglund, P., Storå, J., Götherström, A. & Jakobsson, M. Accurate sex identification of ancient human remains using DNA shotgun sequencing. JAS 40, 4477–4482 (2013).

41 Li, H. et al. The Sequence Alignment/Map format and SAMtools. Bioinformatics 25, 2078–2079 (2009).

42 Devlin, B. & Roeder, K. Genomic control for association studies. Biometrics 55, 997–1004 (1999).

43 Norton, H. L. et al. Genetic evidence for the convergent evolution of light skin in Europeans and East Asians. Mol. Biol. Evol. 24, 710–722 (2007).

44 Bokor, S. et al. Single nucleotide polymorphisms in the FADS gene cluster are associated with delta-5 and delta-6 desaturase activities estimated by serum fatty acid ratios. J. Lipid Res. 51, 2325–2333 (2010).

45 Tanaka, T. et al. Genome-wide association study of plasma polyunsaturated fatty acids in the InCHIANTI Study. PLoS Genet. 5, e1000338 (2009).

46 Uciechowski, P. et al. Susceptibility to tuberculosis is associated with TLR1 polymorphisms resulting in a lack of TLR1 cell surface expression. J. Leukoc. Biol. 90, 377–388 (2011).

47 Wong, S. H. et al. Leprosy and the adaptation of human toll-like receptor 1. PLoSPathog. 6, e1000979 (2010).

48 Ahn, J. et al. Genome-wide association study of circulating vitamin D levels. Hum. Mol. Genet. 19, 2739–2745 (2010).

49 Grundemann, D. et al. Discovery of the ergothioneine transporter. Proc. Natl. Acad. Sci. U. S. A. 102, 5256–5261 (2005).

50 Chauhan, S. et al. ZKSCAN3 is a master transcriptional repressor of autophagy. Mol. Cell 50, 16–28 (2013).

51 Soler Artigas, M. et al. Genome-wide association and large-scale follow up identifies 16 new loci influencing lung function. Nat. Genet. 43, 1082–1090 (2011).

52 Sturm, R. A. et al. A single SNP in an evolutionary conserved region within intron 86 of the HERC2 gene determines human blue-brown eye color. Am. J. Hum. Genet. 82, 424–431 (2008).

53 Eiberg, H. et al. Blue eye color in humans may be caused by a perfectly associated founder mutation in a regulatory element located within the HERC2 gene inhibiting OCA2 expression. Hum. Genet. 123, 177–187 (2008).

54 Pruim, R. J. et al. LocusZoom: regional visualization of genome-wide association scan results. Bioinformatics 26, 2336–2337 (2010).

## References

1 Roodenberg, J. in Life and Death in a Prehistoric Settlement in Northwest Anatolia. The Ilipinar Excavations, Volume III (eds J. Roodenberg & S. Alpaslan Roodenberg) (The Netherlands Institute for the Near East, 2008).

2 Alpaslan Roodenberg, M. S., Todorova, N. & Petrova, V. The Human Burials of Yabalkovo. Praehistorische Zeitschrift 88, 23–37 (2013).

3 Andrews, P., Molleson, T. & Boz, B. in Inhabiting Çatalhöyük: reports from the 1995-1999 seasons. (ed I. Hodder) (British Institute of Archaeology at Ankara, 2007).

4 Karul, N. & Avci, B. Neolithic Communities in the Eastern Marmara: Aktopraklik C. Anatolica 37, 1–15 (2011).

5 Alpaslan Roodenberg, S. in Life and Death in a Prehistoric Settlement in Northwest Anatolia. The Ilipinar Excavations, Volume III (eds J. Roodenberg & S. Alpaslan Roodenberg) (The Netherlands Institute for the Near East, 2008).

6 Roodenberg, J., van As, A., Jacobs, L. & Wijnen, M. H. Early Settlement in the Plain of Yenişehir (NW Anatolia). Anatolica 29, 17–59 (2003).

7 Alpaslan-Roodenberg, S., 2001 - Newly found human remains from Menteşe in the Yenişehir Plain: The season of 2000. Anatolica 27: 1–14.

8 Groenhuijzen M, Kluiving S, Gerritsen FA, Künzel M. (2015) Geoarchaeological Research at Barcın Höyük: Implications for the Neolithisation of Northwest Anatolia. Quarternary International 367: 51–61

9 Gerritsen FA, Özbal R, Thissen L (2013) The Earliest Neolithic Levels at Barcın Höyük, Northwestern Turkey. Anatolica 39, 53–92.

10 Weninger B, Clare L, Gerritsen F, Horejs B, Krauß R, Özbal R, Rohling E. (2014) Neolithisation and Rapic Climate Change (6600-6000 cal BC) in the Aegean and Southeast Europe. Documenta Praehistorica 41, 1–31.

11 During BS (2013) Breaking the Bond: Investigating the Neolithic Expansion in Asia Minor in the Seventh Millennium BC. Journal or World Prehistory 26, 75–100.

12 Carbonell E et al. (2014) Sierra de Atapuerca archaeological sites, in: Sala, R. (Ed.): Pleistocene and Holocene hunter-gatherers in Iberia and the Gibraltar Strait: the current archaeological record. Universidad de Burgos / Fundación Atapuerca. Burgos, 534–560.

13 Cáceres I, Lozano M, Saladié P (2007) Evidence for bronze age cannibalism in El Mirador Cave (Sierra de Atapuerca, Burgos, Spain). Am J Phys Anthropol 133, 899–917.

14 Gómez-Sánchez D et al. (2014). Mitochondrial DNA from El Mirador cave (Atapuerca, Spain) reveals the heterogeneity of Chalcolithic populations. PLoS One 9, e105105.

15 Brandt G, et. Al (2013). Ancient DNA reveals key stages in the formation of central European mitochondrial genetic diversity. Science 342, 257–261.

16 Haak W et al. (2015) Massive migration from the steppe was a source for Indo-European languages in Europe. Nature 522, 207–11

17 Fritsch B, Claßen E, Müller U, Dresely V (2011) Die linienbandkeramischen Gräberfelder von Derenburg „Meerenstieg II“ und Halberstadt „Sonntagsfeld“, Lkr. Harz. Jahresschr. Mitteldt. Vorgesch. 92, 25–229.

18 Leinthaler, B., Bogen, C. & Döhle, H.-J. in Archäologie auf der Überholspur. Ausgrabungen an der A38 Vol. 5 Archäologie in Sachsen-Anhalt, Sonderband (eds V. Dresely & H. Meller) 59–82 (Landesamt für Archäologie Sachsen-Anhalt, Halle (Saale), 2006).

19 Koch, F. Die Glockenbecher-und Aunjetitzer Kultur zwischen Benzingerode und Heimburg - Befunde und Funde der Ausgrabungen an der B 6n. Jahresschrift fuer mitteldeutsche Vorgeschichte 93, 187–290 (2009)

20 Autze, T. in Quer-Schnitt. Ausgrabungen an der B 6n. Benzingerode-Heimburg Vol. 2 Archäologie in Sachsen-Anhalt (eds V. Dresely & H. Meller) 39–51 (Landesamt für Archäologie Sachsen-Anhalt, Halle (Saale), 2005).

21 Dalidowski, X. in Archäologie XXL. Archäologie an der B 6n im Landkreis Quedlinburg Vol. 4 (eds V. Dresely & H. Meller) 116–120 (Landesamt für Archäologie Sachsen-Anhalt, Halle (Saale), 2006).

22 Der Sarkissian, C. et al. Ancient DNA Reveals Prehistoric Gene-Flow From Siberia in the Complex Human Population History of North East Europe. PLoS Genetics 9, e1003296 (2013).

23 Anthony, D. W. (2007) The Horse the Wheel and Language. Princeton Univ. Press, Princeton; Agapov, S.A., I.B. Vasiliev, and V.I. Pestrikova (1990) Khvalynskii Eneoliticheskii Mogil’nik, Saratovskogo Universiteta, Saratov.

24 Vasiliev, I.B., P.F. Kuznetsov, and A.P. Semenova (1994) Potapovskii Kurgannyi Mogil’nik Indoiranskikh Plemen na Volge, Samarskii Universitet, Samara.

## References

1. Allentoft, M. E. et al. Population genomics of Bronze Age Eurasia. Nature 522, 167–172, (2015).

2. Fu, Q. et al. An early modern human from Romania with a recent Neanderthal ancestor. Nature 524, 216–219, (2015).

3. Gamba, C. et al. Genome flux and stasis in a five millennium transect of European prehistory. Nat. Commun. 5, 5257 (2014).

4. Günther, T. et al. Ancient genomes link early farmers from Atapuerca in Spain to modern-day Basques. Proceedings of the National Academy of Sciences, (2015).

5. Haak, W. et al. Massive migration from the steppe was a source for Indo-European languages in Europe. Nature 522, 207–211, (2015).

6. Keller, A. et al. New insights into the Tyrolean Iceman’s origin and phenotype as inferred by whole-genome sequencing. Nat. Commun. 3, 698, (2012).

7. Lazaridis, I. et al. Ancient human genomes suggest three ancestral populations for present-day Europeans. Nature 513, 409–413, (2014).

8. Olalde, I. et al. Derived immune and ancestral pigmentation alleles in a 7,000-year-old Mesolithic European. Nature 507, 225–228, (2014).

9. Olalde, I. et al. A common genetic origin for early farmers from Mediterranean Cardial and Central European LBK cultures. Mol. Biol. Evol., (2015).

10. Raghavan, M. et al. Upper Palaeolithic Siberian genome reveals dual ancestry of Native Americans. Nature 505, 87–91, (2014).

11. Sánchez-Quinto, F. et al. Genomic Affinities of Two 7,000-Year-0ld Iberian Hunter-Gatherers. Curr. Biol. 22, 1494–1499, (2012).

12. Seguin-Orlando, A. et al. Genomic structure in Europeans dating back at least 36,200 years. Science 346, 1113–1118, (2014).

13. Sikora, M. et al. Population Genomic Analysis of Ancient and Modern Genomes Yields New Insights into the Genetic Ancestry of the Tyrolean Iceman and the Genetic Structure of Europe. PLoS Genet. 10, e1004353, (2014).

14. Skoglund, P. et al. Genomic Diversity and Admixture Differs for Stone-Age Scandinavian Foragers and Farmers. Science 344, 747–750, (2014).

15. Skoglund, P. et al. Origins and genetic legacy of Neolithic farmers and hunter-gatherers in Europe. Science 336, 466–469, (2012).

16. Lipson, M. et al. Efficient moment-based inference of admixture parameters and sources of gene flow. Mol. Biol. Evol. 30, 1788–1802, (2013).

17. Patterson, N. et al. Ancient admixture in human history. Genetics 192, 1065–1093, (2012).

18. Bollongino, R. et al. 2000 Years of Parallel Societies in Stone Age Central Europe. Science 342, 479–481, (2013).

19. Bellwood, P. First Farmers: The Origins of Agricultural Societies. (Wiley-Blackwell, 2004).

20. Haak, W. et al. Ancient DNA from European early Neolithic farmers reveals their Near Eastern affinities. PLoS Biol. 8, e1000536, (2010).

21. Brandt, G. et al. Ancient DNA reveals key stages in the formation of central European mitochondrial genetic diversity. Science 342, 257–261, (2013).

22. Szécsényi-Nagy, A. et al. Tracing the genetic origin of Europe’s first farmers reveals insights into their social organization. Proceedings of the Royal Society of London B: Biological Sciences 282, (2015).

23. Underhill, P. A. et al. The phylogenetic and geographic structure of Y-chromosome haplogroup R1a. Eur. J. Hum. Genet. 23, 124–131, (2014).

24. Hollard, C. et al. Strong genetic admixture in the Altai at the Middle Bronze Age revealed by uniparental and ancestry informative markers. Forensic Science International: Genetics 12, 199–207, (2014).

